# Spontaneous oxycodone withdrawal disrupts sleep, circadian, and electrophysiological dynamics in rats

**DOI:** 10.1101/2024.01.21.576572

**Authors:** M Gulledge, WA Carlezon, R Kathryn McHugh, M. Prerau, EH Chartoff

## Abstract

Opioid dependence is defined by an aversive withdrawal syndrome upon drug cessation that can motivate continued drug-taking, development of opioid use disorder, and precipitate relapse. An understudied but common opioid withdrawal symptom is disrupted sleep, reported as both insomnia and daytime sleepiness. Despite the prevalence and severity of sleep disturbances during opioid withdrawal, there is a gap in our understanding of their interactions. The goal of this study was to establish an in-depth, temporal signature of spontaneous oxycodone withdrawal effects on the circadian composition of discrete sleep stages and the dynamic spectral properties of the electroencephalogram (EEG) signal in male rats. We continuously recorded EEG and electromyography (EMG) signals for 8 d of spontaneous withdrawal after a 14-d escalating-dose oxycodone regimen (0.5 - 8.0 mg/kg, 2×d; SC). During withdrawal, there was a profound loss and gradual return of circadian structure in sleep, body temperature, and locomotor activity, as well as increased sleep and wake fragmentation dependent on lights on/off. Withdrawal was associated with significant alterations in the slope of the aperiodic 1/f component of the EEG power spectrum, an established biomarker of arousal level. Early in withdrawal, NREM exhibited an acute flattening and return to baseline of both low (1-4 Hz) and high (15-50 Hz) frequency components of the 1/f spectrum. These findings suggest temporally dependent withdrawal effects on sleep, reflecting the complex way in which the allostatic forces of opioid withdrawal impinge upon sleep and circadian processes. These foundational data based on continuous tracking of nocturnal rhythms, sleep stage composition, and spectral EEG properties provide a detailed construct with which to form and test hypotheses on the mechanisms of opioid-sleep interactions.

## Introduction

Prescription opioid painkillers such as oxycodone exert both analgesic and euphoric effects, the latter of which contribute to high rates of misuse, opioid dependence, opioid use disorder (OUD), and overdose [1]. OUD is a chronic, relapsing disorder characterized by escalating, compulsive drug use despite harmful consequences, a constellation of aversive withdrawal signs, cravings, and relapse [2]. A primary barrier to treatment of OUD is the experience of opioid withdrawal, comprising both acute somatic and protracted affective components [3, 4, 5]. Understanding the spectrum of withdrawal signs and their underlying mechanisms is essential to developing effective treatments.

Sleep disturbances are some of the most highly reported, but least studied, withdrawal symptoms for a number of drugs of abuse, including opioids [6, 7, 8]. Withdrawal-induced sleep disturbances typically include insomnia, fragmented sleep, and excessive daytime sleepiness [6, 9, 10]. Although the specific influence of withdrawal-related sleep disturbance on outcomes remains understudied, sleep disturbance more broadly is associated with increased opioid craving, greater clinical severity, and misuse of benzodiazepines [11, 12, 13]. Indeed, the majority of people with OUD entering outpatient treatment report some sleep disturbance [14]. Although several preclinical and clinical studies have assessed effects of acute opioids on sleep [15, 16, 17], there are relatively few studies characterizing the effects of opioid withdrawal on sleep [8, 18, 19, 20] Notably, two recent papers report disruptive effects of morphine withdrawal on sleep in mice, a nocturnal species. Both studies show increased sleep and decreased wake during lights-off, consistent with the clinical observation of increased daytime sleepiness [19, 20].

Oxycodone activates mu opioid receptors (MOR), which are expressed throughout the brain, including regions that regulate reward, motivation and arousal [18, 21, 22]. Acute MOR activation typically decreases neuronal activity [23], whereas chronic MOR activation results in time-dependent molecular, cellular, and circuit adaptations to counter persistent neuronal inhibition [23, 24, 25, 26]. Upon removal of the opioid (i.e., withdrawal), there is an “unmasking” of these previously MOR-suppressed homeostatic changes, resulting in rebound increases in neuronal activity within MOR-expressing cells [23, 24, 26]. Putative interactions between opioid withdrawal and sleep depend, in part, on the neuroanatomical expression of MORs and their homeostatic responses to chronic opioid exposure and withdrawal. Opioid dependence and withdrawal constitute strong internal and external stimuli that modulate circadian processes [27, 28], homeostatic processes that track sleep need [29, 30, 31, 32], and allostatic processes such as stress [33]. As such, there are numerous possibilities through which opioid withdrawal can influence sleep-wake regulatory networks.

Systems-level neural activity during sleep is commonly measured through the electroencephalogram (EEG), which represents the aggregate dynamics of numerous cortical and subcortical networks over time [34, 35]. Sleep architecture in rodents is categorized into wake, rapid eye movement (REM) sleep, and non-REM (NREM) sleep based on a standardized combination of behavior and EEG events [34]. Thus, sleep EEG analysis can serve as a means of characterizing the effects of opioid withdrawal on sleep dynamics, as well as specific effects on the neural mechanisms governing sleep processes [34, 36, 37]. Broadly, the EEG signal can be decomposed into periodic (oscillatory) and aperiodic (non-oscillatory) components. Periodic components are derived from synchronized activity within and between different brain networks and generally manifest as salient spectral peaks within canonical frequency bands: delta (<4 Hz), theta (4– 8 Hz), alpha (8–12 Hz), beta (12–30 Hz), and gamma (>30 Hz) [37]). In contrast, it is thought that non-oscillatory (aperiodic) components of the power spectrum density (PSD) do not originate from any regular, rhythmic process, but rather from the integration of synaptic currents [38].

While the structure of the aperiodic component is complex [39, 40], it is typically described in terms of a simplified single, or piecewise, power-law model (1/*f*^*α*^), in which spectral power decays as the frequency (*f*) increases [41]. The time constant of decay, *α*, is commonly interpreted as the slope of the linearized log-log transformed EEG power spectrum. Piecewise models will fit this constant separately in high and low frequency components, which are separated by a pivot point (at ∼20-30 Hz in humans) often termed a “knee” [40, 42]. Although the underlying neural mechanisms responsible for the aperiodic component of the EEG are not fully understood, changes in aperiodic slope have been correlated with excitatory-inhibitory (EI) synaptic balance [40, 43]. Increasing evidence shows that pharmacological excitation of neural activity is associated with flatter aperiodic slopes, whereas inhibition is associated with steeper slopes [42, 44]. As such, tracking the aperiodic slope throughout oxycodone withdrawal may provide a temporal signature of changes in global neural activity and arousability.

In this study, male rats were implanted with telemetry devices that wirelessly and continuously emit EEG, EMG, temperature, and activity signals to allow a comprehensive and temporal analysis of how spontaneous oxycodone withdrawal alters sleep states and their associated EEG spectral dynamics. These foundational studies aim to establish signatures of opioid withdrawal on sleep processes that will engender future mechanistic studies and may ultimately be used to inform development of novel therapeutics targeting sleep disruptions.

## Results

### Experimental Overview

To track the effects of spontaneous withdrawal from chronic oxycodone administration on sleep/wake, rats were subcutaneously implanted with telemetry devices (DSI, Data Sciences International) to capture EEG and EMG, temperature, and activity signals and drug pumps (iPrecio) to deliver saline or oxycodone (**Fig. 1A**). Pumps were programmed to deliver drug twice a day (ZT0-2, lights-on and ZT12-14, lights-off). Telemetry data was continuously collected throughout the experiment. The telemetry data from the last 3 days of saline administration (**Fig. 1A**, blue chevron) were used as baseline. Escalating-dose oxycodone was delivered for 14 days, with the last infusion occurring from ZT0-2 on withdrawal day 1 (W1). This allowed testing for naloxone-precipitated withdrawal 30-45 minutes after the last oxycodone infusion ended at ZT2. This study focuses primarily on the temporal effects of spontaneous oxycodone withdrawal measured over withdrawal days 2-8 (W2-W8), (**Fig. 1A**, red chevron). Averaged % time in 3-h bins over 24-h periods in each sleep stage (NREM, REM, Wake; **Fig 1B**) is shown for each of the 4 stages of the experiment: baseline (B, blue), oxycodone (purple), W1 (naloxone, yellow; naloxone, orange), and spontaneous withdrawal W2-W8 (red). Oxycodone dependence is demonstrated on W1 by measuring naloxone-precipitated somatic withdrawal behaviors (**Fig. 1C**).

**Fig 1:**
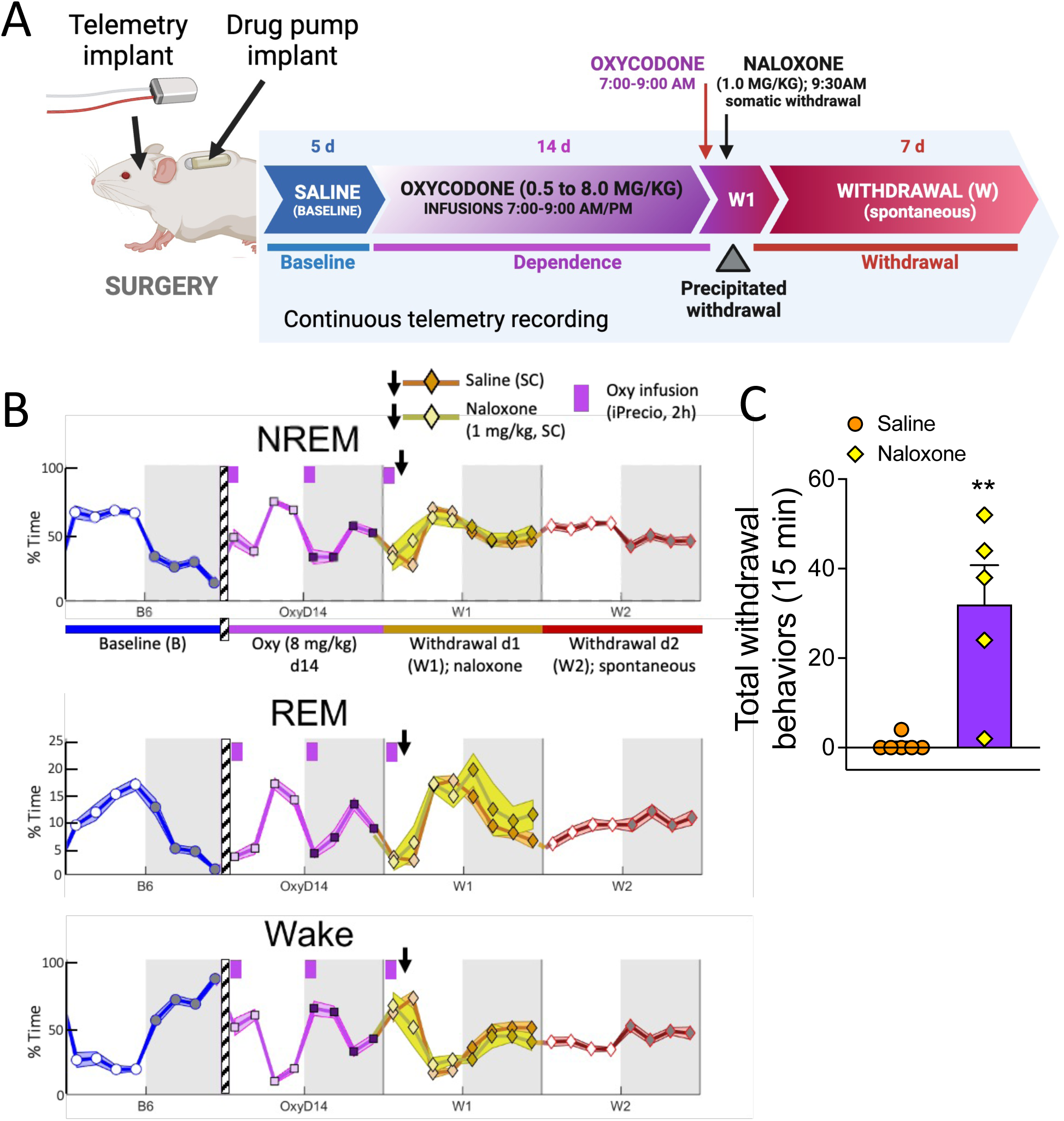
Escalating oxycodone dose infusion protocol modulates sleep stages and produces dependence in male rats. (**A**) Experimental schematic. Rats were implanted subcutaneously (SC) with both a telemetry device and a programmable infusion pump. After recovery, pumps infused saline for 5 days, the last 3 of which were used as baseline sleep recordings. Following saline infusions, pumps delivered escalating-dose oxycodone twice a day (ZT0-2 and ZT12-14) for 14 days (0.5 - 8.0 mg/kg/inf, 2 inf/d). The last infusion occurred from ZT0-2 on withdrawal day 1 (W1). Immediately after the last infusion, rats were injected with either saline or naloxone (1.0 mg/kg, SC). (**B**) % time in each sleep stage (NREM, REM, Wake) presented as average % time ∓ SEM in 3-h bins for baseline day 6 (B6), oxycodone day 14 (8 mg/kg; OxyD14), withdrawal day 1 (W1) divided into saline- (orange diamonds) or naloxone- (yellow diamonds) treated data, and withdrawal day 2 (W2). (**C**) Somatic withdrawal scores from oxycodone-dependent rats on W1, quantified from video recordings 20-35 minutes post-naloxone (yellow diamonds) or saline (orange circles) injections. Total withdrawal behaviors are the sum of ptosis, flattened posture, and stereotyped head bobs instances in the 15-min period. **p<0.001, unpaired t-test, comparing oxycodone-dependent rats injected with saline (N=6) (N=6) or naloxone (N=5). Underlying data is in **“Fig1_Data”**.

### Spontaneous oxycodone withdrawal transiently flattens sleep-wake rhythms

Rodents have a nocturnal sleep/wake rhythm in which most of their sleep (NREM and REM) occurs during the day (lights-on = ZT0-12; subjective day), and most of their Wake occurs during the night (lights-off = ZT12-24; subjective night) [45, 46]. This typical nocturnal sleep/wake rhythm is observed at baseline (B; **Fig. 1B**, blue), whereas the ZT0-2 oxycodone infusion on Oxy d14 suppresses sleep (NREM, REM) and increases Wake for the first 6 hours of lights on (ZT0-6) (**Fig. 1B**, purple). Oxycodone infusions from ZT12-14 (lights off) do not themselves alter % time in sleep stage relative to B, but rather the offset of oxycodone effects (ZT18-24) is associated with increased sleep and decreased wake (**Fig. 1B**, purple). It is important to highlight that telemetry data from each oxycodone infusion over the course of the 14-d escalating dose regimen have been recorded, but are not the focus of this paper. Here we are specifically interested in how oxycodone withdrawal impacts sleep.

To characterize the effects of spontaneous oxycodone withdrawal on the circadian rhythmicity of sleep architecture observed during baseline recording days, we plotted the percent of total time spent in NREM, REM, and Wake (**Fig. 2A, B, C**, respectively) in 3-h bins covering the last 3 days of baseline recordings (saline infusions, B4, B5, B6) and spontaneous oxycodone withdrawal days 2-8 (W2-W8). Withdrawal day 1 (W1) is not included in our spontaneous withdrawal results because rats received their last 2-h infusion of oxycodone and an injection of either naloxone or saline on the morning of W1 (see Methods). For all sleep states, we observe typical circadian rhythmicity during B4-B6, followed by a marked flattening (loss of rhythmicity) during early withdrawal (∼W2-W4; **Fig. 2A, B, C**) and a gradual return towards baseline patterns by W8. To quantify the effects of spontaneous oxycodone withdrawal on circadian-like rhythmicity during NREM, REM, and Wake, we compared the magnitude and timing of several sleep stage features during W2-8 to the average of the last 3 baseline days (BL_avg_). First, we computed the circadian index (CI), defined here as the peak to trough amplitude in the 3-h bin values on a particular day, normalized to each individual rat’s average peak to trough amplitude over the last 3 baseline days (BL_avg_). A decrease in CI relative to BL_avg_ indicates a reduction in the magnitude of circadian rhythmicity (i.e., flattening). For the % of time in each sleep state, Friedman’s Tests show that the CI is significantly reduced compared to BL_avg_: [NREM, Q = 44.98, p<0.0001, **Fig. 2D**, left; REM, Q = 31.82, p<0.0001, **Fig. 2E**, left; Wake, Q = 45.18, p<0.0001, **Fig. 2F**, left).

**Fig 2.**
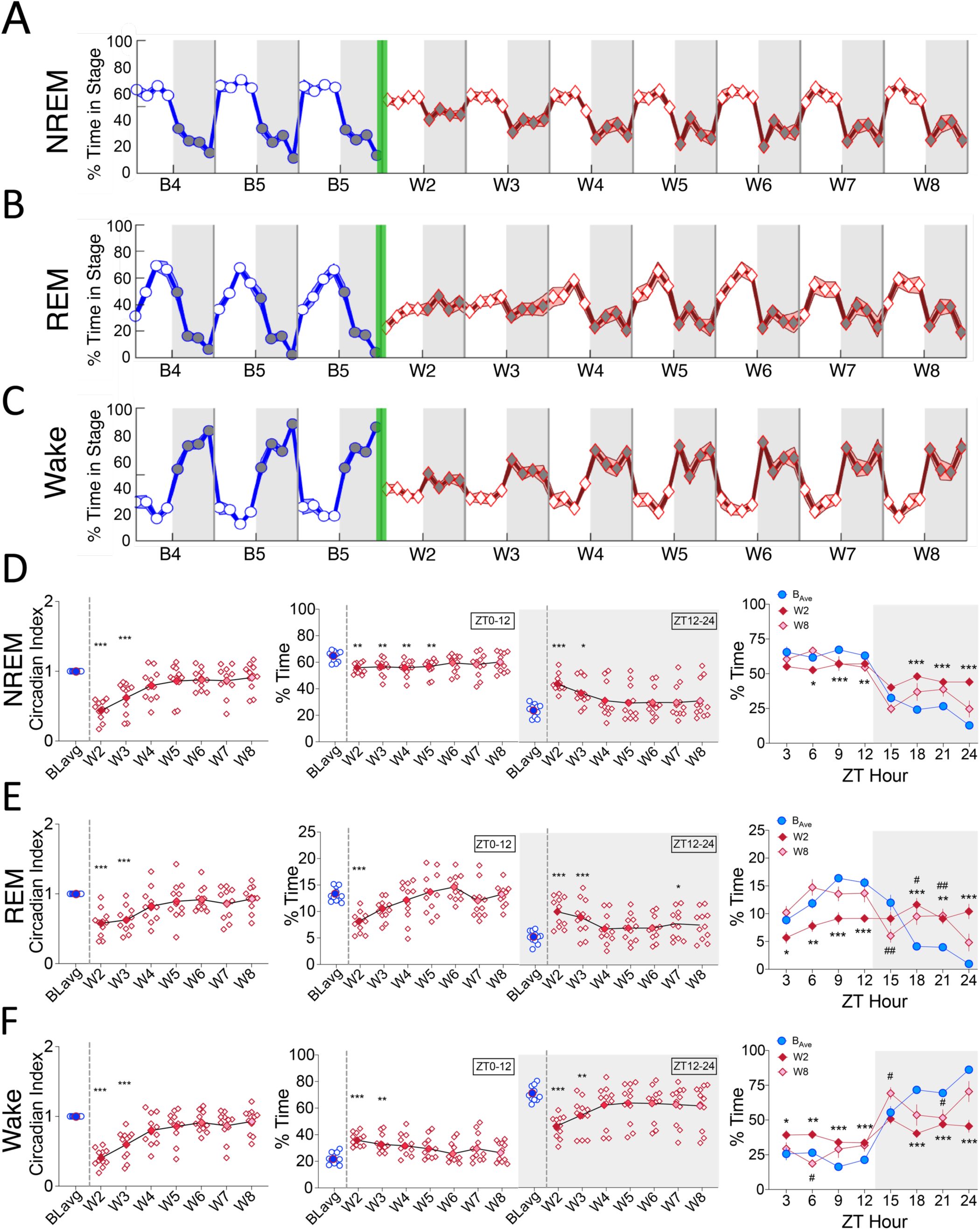
Spontaneous oxycodone withdrawal transiently flattens circadian sleep rhythms. The mean % of total time in 3-h bins (∓SEM) spent in each sleep stage [(**A**) NREM, (**B**) REM, and (**C**)Wake] is shown for baseline days 4, 5, and 6 (B4, B5, B6; blue line, circles) and spontaneous withdrawal days 2-8 (W2-W8; red line, diamonds). Vertical green bars between B6 and W2 represent the 14-d escalating oxycodone dose regimen and W1. Grey shaded regions represent ZT12-24 (lights off), and unshaded regions represent ZT0-12 (lights on). For NREM, REM, and Wake (**D, E, F**, respectively), the mean Circadian Index (CI; left panels), % total time (ZT0-12 or ZT12-24; center panels), and the mean % of total time in 3-h bins for BL_avg_, W2, and W8 (right panels) are shown. CI is defined as each days’ maximum - minimum 3-h bin difference divided by the individual rat’s average baseline difference. For D, E, F, left and middle panels: *p<0.05, **p<0.01, ***p<0.001, Dunn’s post hoc tests. For D, E, F, right panels, *p<0.05, **p<0.01, ***p<0.001 comparing W2 to BLavg, and #p<0.05, ##p<0.01 comparing W8 to BLavg, Dunnett’s post hoc tests. N=11 rats. Underlying data is in **Fig2_Data**. Abbreviations: ZT, zeitgeber time; BL_avg_, average of baseline days 4, 5, 6; W, withdrawal.

To determine whether the effects of oxycodone withdrawal on sleep/wake occur primarily during lights on or off, we quantified the average percent time spent in NREM, REM and Wake for ZT0-12 and for ZT12-24 (**Fig 2D, E, F**, middle panels) and compared W2-W8 to BL_avg_. Overall, we find that during early withdrawal the % time asleep decreases, whereas the % time awake increases, during lights on (NREM, Q = 22.24, p < 0.005, **Fig. 2D**, middle; REM, Q = 40.55, p<0.0001, **Fig. 2E**, middle; Wake, Q = 28.92, p < 0.0001, **Fig. 2F**, middle). In contrast, there is a general increase in % time asleep and decrease in % time awake during lights off (NREM, Q = 32.86, p<0.0001, **Fig. 2D**, middle gray; REM, Q = 30.18, p<0.0001, **Fig. 2E**, middle gray; Wake, Q = 33.36, p<0.0001, **Fig. 2F**, middle gray). Together with the CI data, these findings indicate that the most robust effects of spontaneous withdrawal from 14 days of chronic, escalating-dose oxycodone occur during W2 and W3. Further, there is an association between loss of circadian rhythmicity and insomnia-like reductions in time asleep and subsequent increases in lights off (active period) time asleep.

Thus far, our data suggest a robust flattening of circadian-like rhythmicity for each sleep stage on W2 with day-by-day improvements toward baseline. We took a more granular look at the % time in each sleep stage on W8 compared to BL_avg_ and W2 by examining the data in 3-h bins (**Fig. 2D, E, F**; right panels). As expected, the % time in each sleep stage on W2 is relatively flat from ZT0-ZT24, with little difference between lights on and off. On W8; however, recovery towards the rhythmicity of BL_avg_ is observed during lights on, but not lights off. Indeed, similar to W2, the % time in REM and Wake on W8 during several 3-h bins in lights off is significantly different from BL_avg_. These results depend on significant interactions (Treatment day × ZT hour) from 2-way repeated measures ANOVA tests followed by Dunnett’s multiple comparison tests: REM, F_(4.895, 48.95)_=13.13, p<0.0001, **Fig. 2E**, right panel; Wake, F_(3.330, 33.30)_=11.44, p<0.0001, **Fig. 2F**, right panel.

Similar to effects on sleep architecture rhythmicity, spontaneous oxycodone withdrawal reduces the magnitude of activity and temperature circadian rhythmicity (**Fig. 3A, B**, respectively), with the greatest effects occurring on W2 and W3 and subsequent recovery towards BLavg by W8. For temperature and activity, Friedman’s Tests show that the CI is significantly reduced compared to BL_avg_: Temp, Q = 34.85, p<0.0001, **Fig. 3C**, left panel; Activity, Q = 48.52, p<0.0001, **Fig. 3D**, left panel. In contrast to the effects of oxycodone withdrawal on sleep; effects on activity and temperature rhythmicity occurred primarily during lights off. This is particularly evident in **Fig. 3C, D**, right panels, in which 2-way repeated measures ANOVAs show significant Treatment day × ZT hour interactions (Temp, F_(3.293, 32.93)_ = 7.381, p<0.001, **Fig. 3C**, right panel; Activity, F_(2.742, 27.42)_ = 7.953, p<0.001, **Fig. 3D**, right panel), and Dunnett’s post hoc tests show that significant pairwise comparisons occur during lights off.

**Fig 3.**
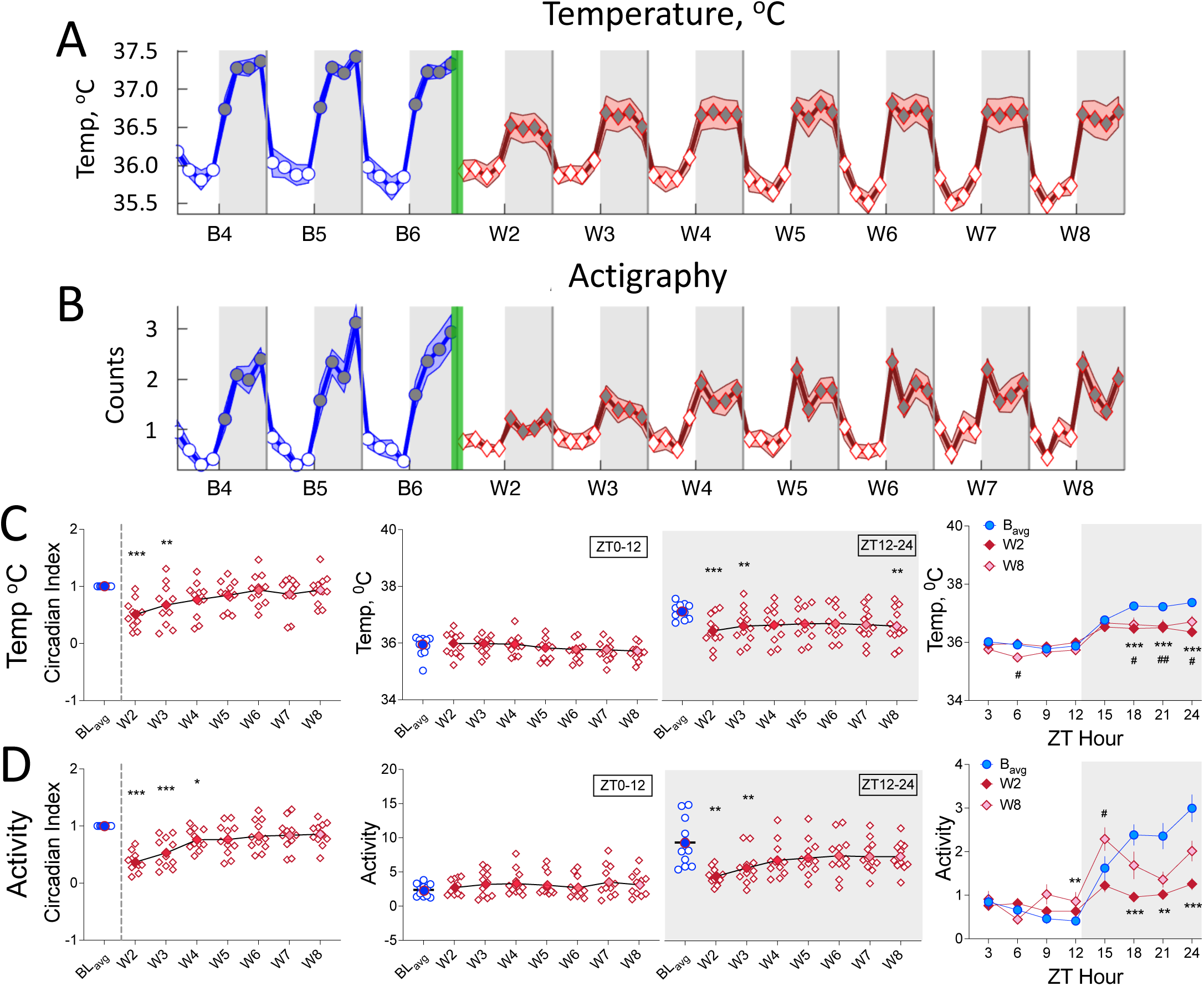
Spontaneous oxycodone withdrawal transiently reduces temperature and activity during lights off. The mean temperature (°C; **A**) and activity counts (in thousands) (**B**) in 3-h bins (∓SEM) are shown for baseline days 4, 5, and 6 (B4, B5, B6; blue line, circles) and for spontaneous withdrawal days 2-8 (W2-W8; red line, diamonds). Vertical green bars between B6 and W2 represent the 14-d escalating oxycodone dose regimen and W1. Grey shaded regions represent ZT12-24 (lights off), and unshaded regions represent ZT0-12 (lights on). For temperature and activity (**C, D**, respectively), Circadian Index (CI; **left panels**), % total temperature (**C, middle panel**) or activity counts (in hundreds) (**D, middle panel**) during ZT0-12 or ZT12-24, and mean temperature (**C, right panel**) or activity counts (**D, right panel**) in 3-h bins for BL_avg_, W2, and W8 are shown. For **C, D, left and middle panels**: *p<0.05, **p<0.01, ***p<0.001, Dunn’s post hoc tests. For **C, D, right panels**, **p<0.01, ***p<0.001 comparing W2 to BL_avg_, and #p<0.05, ##p<0.01 comparing W8 to BL_avg_. N=11 rats. Underlying data is in **Fig3_Data.** *Abbreviations*: ZT, zeitgeber time; BL_avg_, Baseline average; W, withdrawal.

### Both sleep and wake become increasingly fragmented during early spontaneous oxycodone withdrawal

Above, we demonstrate that during early spontaneous withdrawal there are overall reductions in the amplitudes of circadian-like rhythms for each sleep stage, temperature, and activity, which result in overall changes in the amount of sleep/wake during lights on and off. We next investigated whether oxycodone withdrawal affects sleep stage switching, either through increased consolidation or fragmentation of sleep stage bouts. Hypnograms (experimenter-scored sleep stages plotted over time; **Fig. 4A**) are shown for B4-B6 and W2-W8 from a representative oxycodone-treated rat. The hypnograms show marked differences between baseline days and W2, 3. Qualitatively, the long-duration wake bouts predominant during lights-off decrease in length on W2, replaced with an increased number of sleep bouts, suggesting increased sleep stage fragmentation on W2 and W3 relative to baseline. By eye (**Fig. 4A**), this loss of lights-on/lights-off rhythmicity recovers around W4 and continues towards baseline patterns by W8.

**Fig. 4.**
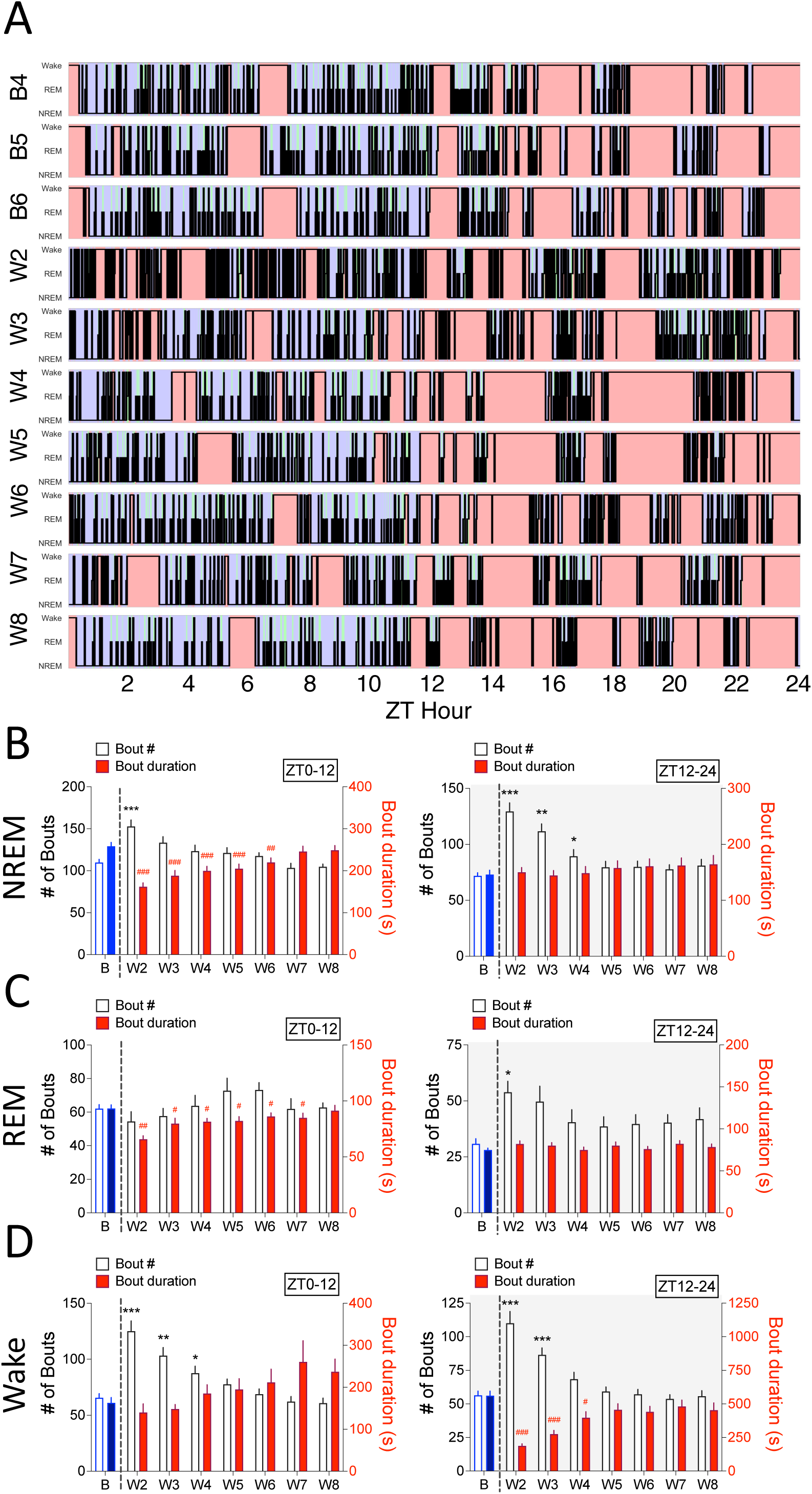
Spontaneous oxycodone withdrawal initially increases sleep fragmentation. (**A**) Hypnogram data from a representative oxycodone-treated rat is plotted for each of B4-B6 and W2-W8 days, with Wake represented at the top of each hypnogram with red shading, REM in the middle with green shading and NREM at the bottom with blue shading. For NREM, REM, and Wake (**B, C, D**, respectively) the mean (∓SEM) number of bouts (white bars, left Y axis) and bout durations (filled bars, right Y axis) are shown for BL_avg_ and W2-W8 for ZT0-12 (**left panels**) and ZT12-24 (**right panels**). For lights on, when rats show less % time asleep, the number of NREM (**B, left panel**) and Wake (**D, left panel**) bouts is increased, with corresponding decreases in NREM and REM bout duration. For lights off, there is a significant increase in the # of bouts of all sleep stages (**B, C, D, right panels**) with a corresponding decrease in Wake bout duration (**D, right panel**). *p<0.05, **p<0.01, ***p<0.001, comparing bout # during withdrawal to BL_avg_ bout #, Dunnett’s post hoc tests. #p<0.05, ##p<0.01, ###p<0.001 comparing bout duration during withdrawal to BL_avg_ bout duration, Dunnett’s post hoc tests. N=11 rats. Underlying data is in **Fig4_Data.** *Abbreviations*: ZT, zeitgeber time; Baseline (B4, B5, B6 average), BL_avg_; W, withdrawal.

To characterize the degree to which spontaneous oxycodone withdrawal alters sleep/wake fragmentation, we quantified bout number and bout length (s) for each sleep stage during either lights-on (**Fig. 4B, C, D**; left panels) or lights-off (**Fig 4B, C, D**; right panels) and compared these to their respective BL_avgs_. For any given arousal stage (NREM, REM, Wake), an increase in the number of bouts with a concurrent decrease - or no change in - bout duration indicates fragmentation [47, 48]. Consistent with what is clear to the eye from **Fig 4A**, the most robust effect is increased Wake fragmentation during both lights on and off (**Fig. 4D**). This is demonstrated with significant Withdrawal Day × Bout Measure interactions (Wake, lights on: F_(1.904, 19.04)_ = 6.075; p<0.01, **Fig4D**, left; Wake, lights off: F_(3.474, 34.74)_ = 29.92, p<0.0001, **Fig4D**, right).

During lights on, the number of Wake bouts increases compared to BL_avg_, but the bout length does not change. This effect requires a decrease in bout number or length in NREM and/or REM. Two-way repeated measures ANOVAs show that NREM and REM bout lengths decrease: a significant Withdrawal Day × Bout Measure interaction for NREM during lights on (F_(7, 70)_ = 25.71; p<0.0001, **Fig4B**, left) and a main effect of Day for REM during lights on (F_(2.41, 24.10)_ = 11.74; p<0.0001, **Fig. 4C**, left). Interestingly, increased Wake fragmentation ends on W4, yet NREM and REM bout durations remain significantly decreased until W6 and W7, respectively. During lights off, the mean number of Wake bouts dramatically increases from 54.45 (∓3.02 SEM) at BL_avg_ to 110.09 (∓8.72 SEM) on W2 and 86.54 (∓5.08 SEM) on W3, with concurrent decreases in bout length from 561.73s (∓33.34 SEM) at BL_avg_ to 187.18s (∓15.31SEM) on W2 and 275.64s (∓26.92SEM) on W3. The increase in Wake fragmentation during lights off is accompanied by significant increases in the number of NREM and REM bouts, as shown by a Withdrawal Day × Bout Measure interaction (NREM, lights off: F_(2.561, 25.61)_ = 7.524; p<0.005, **Fig. 4B**, right) and a main effect of Day (REM, lights off: F_(3.46, 34.60)_ = 5.681; p<0.005, **Fig. 4C**, right). To summarize, oxycodone withdrawal results in profound arousal and sleep state fragmentation. Wake is the most fragmented - both during lights on and off. NREM also shows significant fragmentation during lights on and off, while W2 during lights off is the only time during which REM sleep shows fragmentation. All of these changes suggest both severe disruption of normal rat sleep architecture during early oxycodone withdrawal and differential effects on the neural circuitry that governs sleep/wake initiation maintenance.

### Spectral power and aperiodic EEG aperiodic slope are differentially and temporally altered during spontaneous oxycodone withdrawal

Beyond sleep architecture and circadian rhythms, it is vital to understand if, and how, spontaneous oxycodone withdrawal affects the magnitude, directionality, and temporal dynamics of functional network activity underlying sleep. To those ends, we examined EEG spectral dynamics for NREM, REM, and Wake across baseline B4-B6 and withdrawal W2-W8 days. We first derived estimates of relative spectral power for the canonical frequency bands delta (1-4 Hz), theta (4–8 Hz), alpha (8–12 Hz), beta (12–30 Hz), and gamma (30-50 Hz). Spontaneous oxycodone withdrawal had the most robust effects on spectral power and aperiodic 1/f slopes during NREM. As such, our results and discussion focus on NREM, although data from REM and Wake is presented in **Supplementary Figures**.

#### Spontaneous oxycodone withdrawal transiently flattens the circadian-like rhythmicity of NREM relative spectral power, but not that of REM or Wake

Given the flattening of baseline circadian rhythms within sleep architecture, we next analyzed EEG relative spectral power to assess any effects observed concurrently during spontaneous oxycodone withdrawal. Qualitatively, NREM relative power measured over baseline (B4-B6) shows a clear rhythmicity [49] for each canonical frequency band (**Fig 5**), with the exception of theta (**Fig. 5B**). For delta, relative power is highest at the beginning of lights-on (ZT0), decreases until ZT12 where it is at its lowest, and increases again within ZT12-24. In contrast, alpha, beta, and gamma show the opposite pattern, with the lowest relative power occuring at ZT0. During withdrawal, NREM rhythms of power appear to flatten before gradually recovering to baseline periodic structure by W8. To determine if the rhythmicity of power was affected by oxycodone withdrawal, we quantified the CI for NREM across all frequency bands (**Fig. 5A-E; left panels**). A Friedman’s Test found significant decreases to the CI for every frequency band; including theta, which had a qualitatively unclear rhythm (Delta, Q = 47.91, p<0.0001; **Fig. 5A**; Theta, Q = 26.39, p<0.0005; **Fig. 5B**; Alpha, Q = 52.45, p<0.0001; **Fig. 5C**; Beta, Q = 54.36, p<0.0001; **Fig. 5D**; Gamma, Q = 40.09, p<0.0001; **Fig. 5E**). Thus, spontaneous oxycodone withdrawal flattens NREM rhythms for all frequency bands.

**Fig 5:**
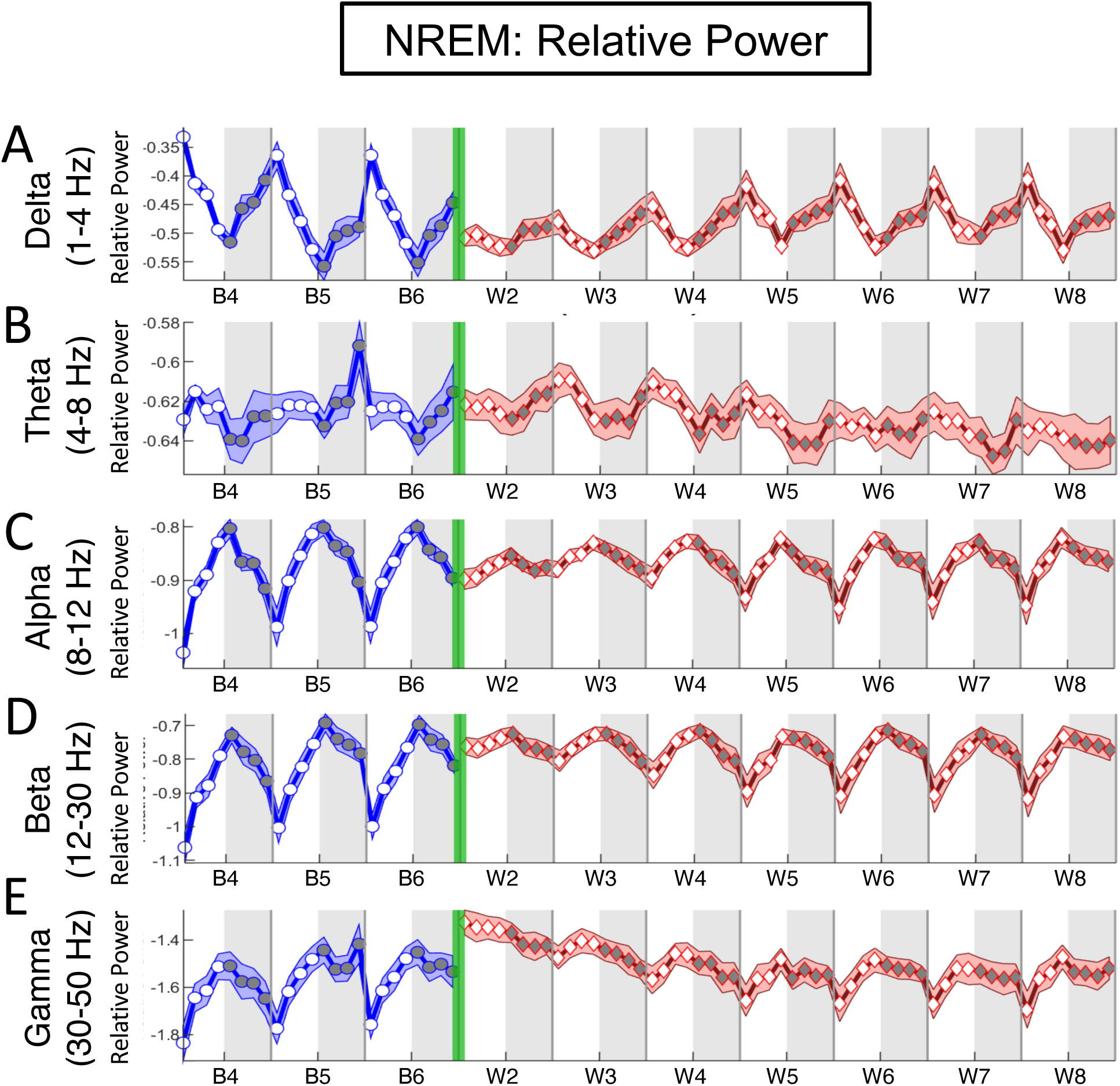
Spontaneous oxycodone withdrawal transiently eliminates circadian-like changes in NREM depth. The mean relative power for NREM is plotted in 3-h bins (∓SEM) for (**A**) Delta (1-4Hz), (**B**) Theta (4-8Hz), (**C**) Alpha (8-12Hz), (**D**) Beta (12-30Hz) and (**E**) Gamma (30-50Hz) frequency bands throughout baseline days 4, 5, and 6 (B4, B5, B6; blue line, circles) and spontaneous withdrawal days 2-8 (W2-W8; red line, diamonds). Vertical green bars between B6 and W2 represent the 14-d escalating oxycodone dose regimen and W1. Grey shaded regions represent ZT12-24 (lights off), and unshaded regions represent ZT0-12 (lights on). Oxycodone withdrawal has a global effect on relative power, rather than modulating specific frequency ranges. N=11 rats. Underlying data is in **Fig5_Data.**

Qualitatively, REM and Wake relative power measured over baseline B4-B6 days appear to show subtle rhythmicities at individual frequency bands (**S1, S3, Figs,** respectively), but more detailed analyses of these potential rhythms were not done, as they are beyond the scope of this foundational study. Furthermore, CI analyses did not reveal any significant differences between relative power at BL_avg_ compared to W2-W8 across frequency bands for either sleep stage (**S2,** REM; **S4,** Wake **Figs, left panels**).

#### Early spontaneous oxycodone withdrawal has global effects on NREM relative power across frequencies, with minimal effects on REM and Wake

We next quantified the mean NREM, REM, and Wake relative power for BL_avg_ and W2-W8 during either lights-on or lights-off **(Fig 6**, NREM; **S2 Fig, middle panels**, REM; **S4, middle panels,** Wake) and performed Friedman’s Tests followed by Dunn’s post hoc tests for significance. For NREM, spontaneous oxycodone withdrawal shows a continuous gradient in power changes with increasing frequency during lights-on, but sparse changes in lights-off. Compared to BL_avg_, relative power during NREM delta is significantly decreased for lights on (Q = 33.12, p<0.0001; **Fig. 6A**, **middle white**). Between 4-12 Hz (theta and alpha) there is a switch in directionality such that relative power in higher frequencies (12-50 Hz, beta and gamma) is significantly increased for lights on (beta, Q = 37.15, p<0.0001; **Fig. 6D**, middle white; gamma, Q = 53.55, p<0.0001; **Fig. 6E**, **middle white**). During lights-off, NREM relative power shows a few significant changes compared to BL_avg_. Interestingly, these (moderate) changes peak on later withdrawal days than those observed for % time in sleep states, sleep fragmentation, and circadian-like rhythmicity. Specifically, the peak increase in NREM relative power compared to BL_avg_ during lights-off occurs on W5 for delta (1-4 Hz; Q = 19.85, p<0.01; p<0.005 at W5, Dunn’s test, **Fig 6A, middle gray panel**) and the peak decrease in relative power occurs on W5 for alpha (8-12 Hz; Q = 20.55, p<0.005; p<0.005 at W5, Dunn’s test, **Fig 6C, middle gray panel**). The strong effect of oxycodone withdrawal on NREM relative power magnitude during lights-on compared to lights-off can best be seen in the right panels of **Fig 6**. In particular, spontaneous oxycodone withdrawal dramatically decreases delta power and increases beta and gamma power during lights-on, with the maximum effects on W2 occurring between ZT0-9. This is demonstrated with 2-way repeated measures ANOVAs and significant 3-h bin × Treatment Day interactions (delta, F_(2.855, 28.55)_ = 13.24, p<0.0001, **Fig 6A, right panel**; beta, F_(2.604, 26.04)_ = 17.46, p<0.0001, **Fig 6D, right panel**; gamma, F_(4.028, 40.28)_ = 19.92, p<0.0001, **Fig 6E, right panel**). This may suggest that oxycodone withdrawal has long-lasting effects on oscillatory activity when transitioning into the sleep period (lights-on). Across W2-W8, oxycodone withdrawal did not have robust or consistent effects on REM or Wake power during lights-on/off, as analyzed and shown in **S2 Fig**, REM and **S4 Fig**, Wake. A finding shared by all three sleep stages is that relative power is at BL_avg_ levels by W8 across all frequency bands, broadly similar to % time in sleep stages and sleep fragmentation (**Figs 2, 4**).

**Fig 6:**
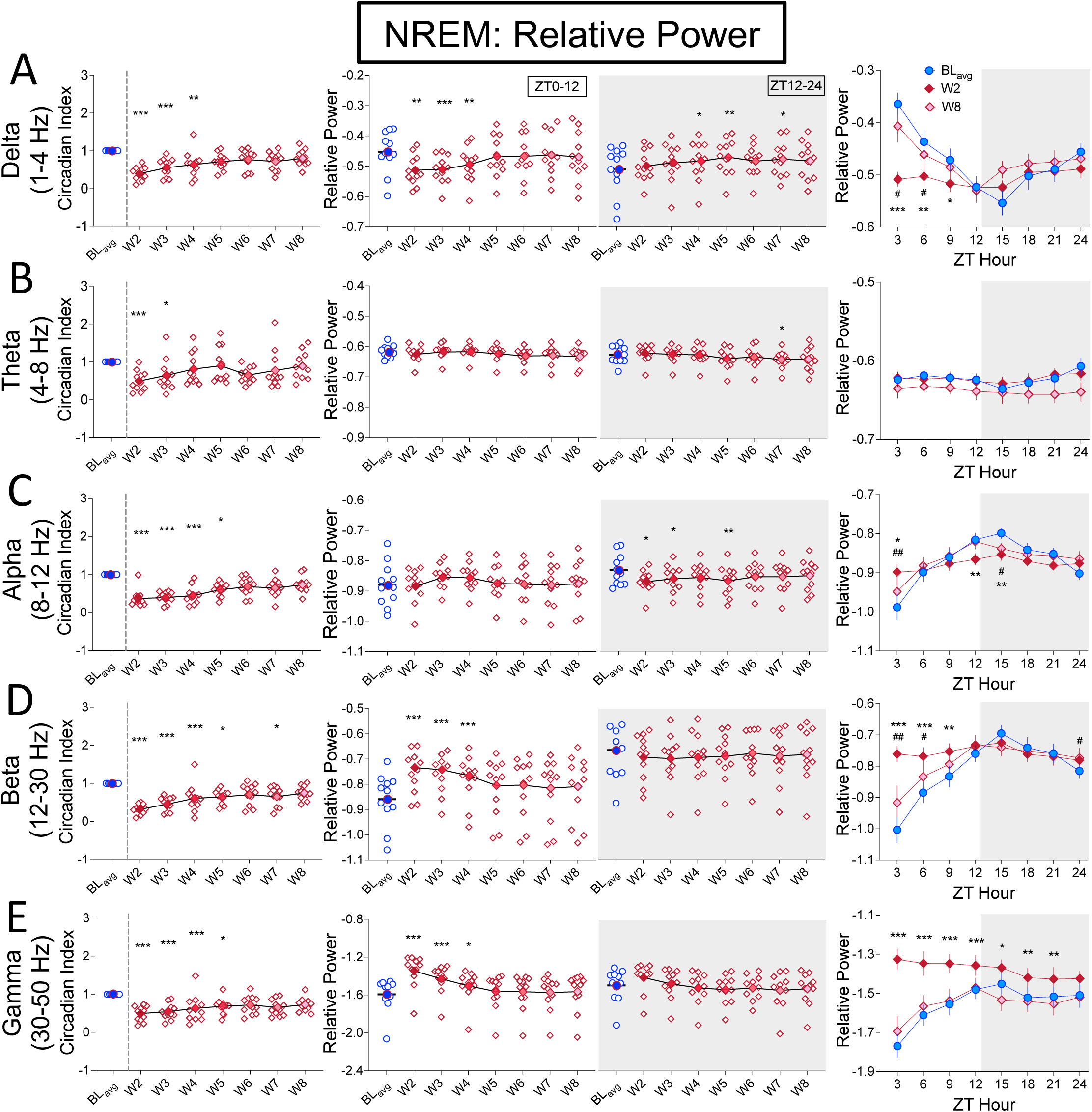
Spontaneous oxycodone withdrawal initially flattens NREM power rhythms, decreases low frequency and increases high frequency power. For relative power during NREM, the mean (∓SEM) Circadian Index (CI; **left panels**), mean (∓SEM) relative power per frequency band for ZT0-12 or ZT12-24; **center panels**, and the mean (∓SEM) relative power per frequency band in 3-h bins for BL_avg_, W2, and W8 (**right panels**) are shown for (**A**) Delta 1-4Hz, (**B**) Theta (4-8Hz), (**C**) Alpha (8-12Hz) (**D**) Beta (12-30Hz), (**E**) Gamma (30-50Hz) frequency bands. Non-parametric Friedman with Dunn’s post hoc tests (**left and center panels**), and two-way repeated measures ANOVA with Dunnett’s post hoc tests (**right panels**) compared W2-W8 to B_avg_. ***p<0.001.**p<0.01, *p<0.05. N=11 rats. Underlying data is in Fig6_Data. *Abbreviations*: ZT, zeitgeber time; BL_avg_, Baseline average; W, withdrawal.

#### Spontaneous oxycodone withdrawal transiently flattens NREM aperiodic slopes at low frequencies, with minimal effects on REM and Wake

The gradient of spectral power changes observed in lights-on NREM during spontaneous withdrawal suggests that, rather than a series of variable band-specific changes, what we are observing may be a global phenomenon across the entire spectrum. Specifically, these changes are consistent with a “tilt” in the aperiodic spectrum around theta/alpha ranges. We therefore explicitly characterize this phenomenon by quantifying the aperiodic piece-wise low frequency (LF, 1-4Hz) and high frequency (HF, 15-50Hz) spectral slope and their changes during the experiment. For visualization, we show example NREM relative spectra (in log-log) taken from a rat at baseline (BL_avg_), W2, and W8 during lights-on (ZT0-12) and during lights-off (ZT12-24) (**Fig 7A, top panels**) with a close-up of the HF and LF regions (**Fig 7A, lower panels)**. For the LF lights-on (lower, left), we observe a distinct flattening of the spectral slope at W2 (red), pivoting down from a point at ∼4Hz. For the HF-lights on and lights off (lower, right), we also see a flattening in the W2 (red) slope, but this time pivoting up from the lower end at ∼15 Hz.

**Fig 7.**
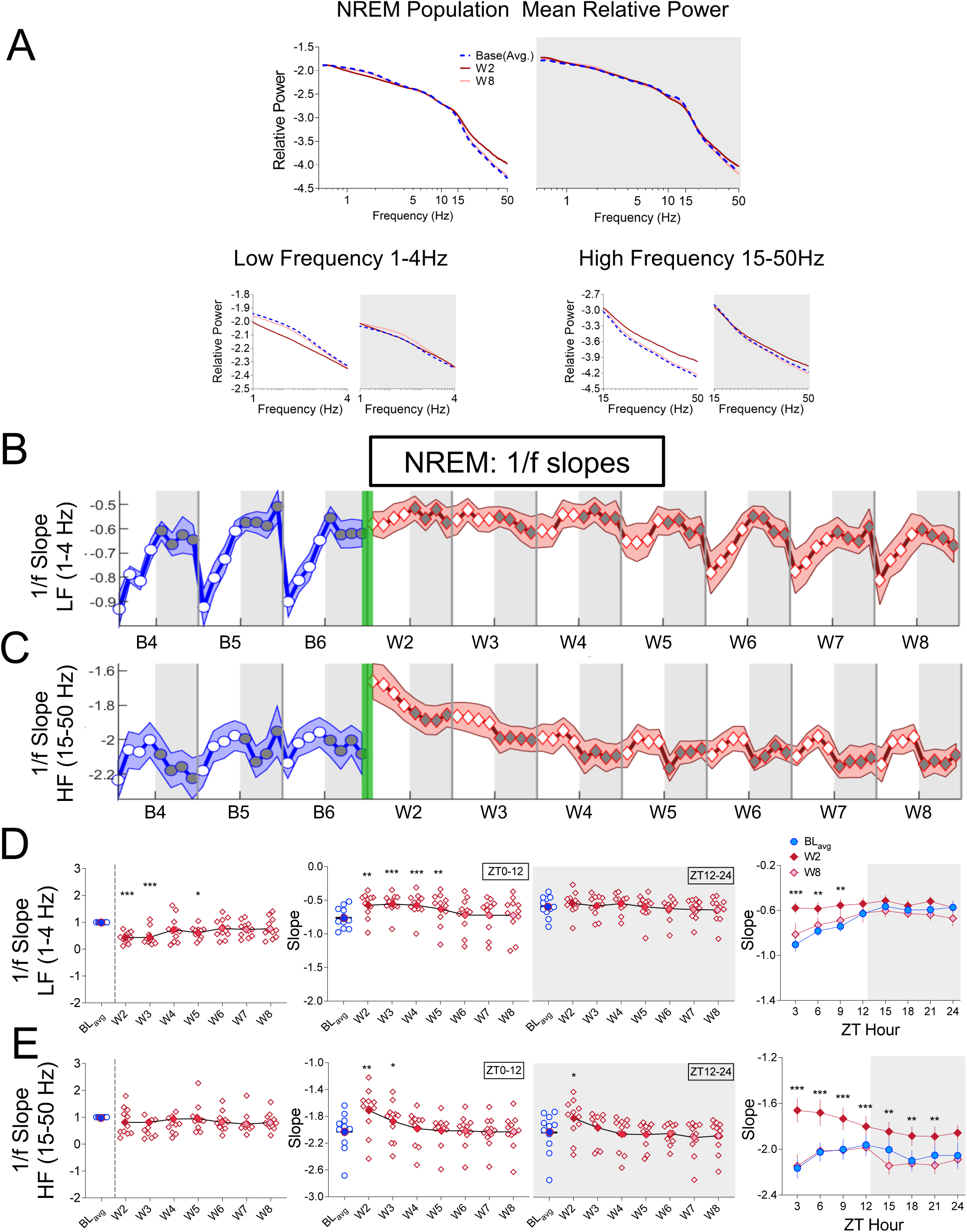
NREM 1/f slopes calculated from Power Spectrum Densities (PSDs) have a robust nocturnal rhythm at low frequencies that are initially flattened during spontaneous oxycodone withdrawal. (**A, top panel**) The average NREM PSD normalized to each rat’s total power from 0.5-50Hz plotted on a log-log scale for lights-on (ZT0-12 unshaded) and lights-off (ZT12-24; shaded) for BL_avg_ (blue), W2 (red), and W8 (pink). (**A, bottom panels**) Zoomed-in low frequency (LF, 1-4 Hz) and high frequency (HF, 15-50 Hz) portions of the NREM PSD during lights-on and lights-off are plotted and used to calculate LF and HF 1/f slopes. The mean NREM 1/f slope (∓SEM) per 3-h bin is plotted for (**B)** low frequency and (**C**) high frequency ranges across baseline days 4, 5, and 6 (B4, B5, B6; blue line, circles) and spontaneous withdrawal days 2-8 (W2-W8; red line, diamonds). Vertical green bars between B6 and W2 represent the 14-d escalating oxycodone regimen and W1. For both LF (**D**) and HF (**E**) NREM 1/f slopes, the mean (∓SEM; including individual rat values) Circadian Index (**left panels**), 1/f slopes (ZT0-12 or ZT12-24; **center panels**), and 3-h bin data for BL_avg_, W2, and W8 (**right panels**) are shown. Non-parametric Friedman tests with Dunn’s post hoc tests (**left and center panels**), and two-way repeated measures ANOVA with Dunnett’s post hoc tests (**right panels**) compared W2-W8 to B_avg_. ***p<0.001.**p<0.01, *p<0.05. N=11 rats. Underlying data is in Fig7_Data. *Abbreviations*: ZT, zeitgeber time; BL_avg_, Baseline average; W, withdrawal.

To characterize aperiodic spectral dynamics baseline and withdrawal days, we computed LF and HF slopes in 3-h bins for each sleep stage (**Fig 7B, C**, NREM; **S5A, B Fig**, REM; **S6A, B Fig**, Wake). As with relative power, we found that the most robust effects of oxycodone withdrawal on aperiodic activity occurred during NREM. Interestingly, the LF aperiodic signal (**Fig 7B**, blue) has a robust, circadian-like rhythm at baseline that is similar in shape and directionality to the NREM periodic alpha, beta and gamma rhythms (**Fig 5C, D, E**). To quantify rhythmicity changes to aperiodic slope at either LF or HF, we calculated the circadian index across all sleep stages by comparing W2-8 to BL_avg_ (**Fig 7D, E,** NREM**; S5C, D Fig,** REM**; S6C, D Fig,** Wake**; left panels**). For NREM, oxycodone withdrawal significantly decreases the LF slope CIs. Friedman’s test: Q = 31.33, p<0.0001, **Fig. 7D**, left). HF slope CIs did not change during withdrawal (**Fig. 7E**, left). REM and Wake slopes appear highly variable and complex (**S5, S6 Figs**), and quantification of slope CIs reveal no significant differences compared to REM or Wake BL_avg_ (**S5C, D,** REM; **S6C**, **D,** Wake, Figs, **left panels**). The one exception is a significant decrease in REM CI for HF slopes only on W3: Friedman’s test, Q=22.42, p<0.005, Dunn’s test, p<0.005 for W3 vs BL_avg_ (**S5D Fig**, REM, **left panel**).

To quantify the daily changes in slopes during withdrawal, we plotted NREM, REM, and Wake LF and HF slopes for BL_avg_ and W2-W8 separately for lights-on and lights-off (**Fig 7D**, **E**, NREM; **S5C, D Fig**, REM; **S6C, D Fig**, Wake; **middle panels**). We found significantly different slope magnitudes during withdrawal for all sleep stages, with the greatest effects in NREM. For NREM, the LF slope is significantly closer to 0 (indicates the slope is becoming flatter) on W2-W5 during lights-on when compared to BL_avg_ (Q = 41.21, p<0.0001; **Fig. 7D**, middle white) but not during lights-off. This may suggest that oxycodone withdrawal prevents normal light induced changes to LF aperiodic activity. The effects of withdrawal on lights-on sleep metrics are the most prolonged for NREM. During withdrawal, NREM HF 1/f slopes are also significantly flattened (approaching 0) during lights on (Q = 44.48, p<0.0001; **Fig. 7E**, middle white), but this effect is only on W2 and W3. In addition, the flattening of NREM HF 1/f slopes on W2 remains significant during lights-off (Q = 36.33, p<0.0001; **Fig. 7E**, middle gray). For REM, slopes are significantly different compared to BLavg during lights-on and lights-off for LF (lights-on: Q = 22.79, p<0.005; **S5C Fig,** middle white; lights-off: Q = 32.80, p<0.0001; **S5 FigC,** middle gray), and during lights-off for HF (lights-off: Q = 25.0, p<0.001; **S5D Fig,** middle gray). Slopes at LF become steeper (albeit slightly), unlike those for NREM. We report these results, but the nominal effects combined with the low percent of total sleep time spent in REM suggest these findings need to be replicated and/or measured more directly before drawing firm conclusions.

Aperiodic activity follows the same patterns as those of power, sleep/wake %, activity, and temperature wherein oxycodone withdrawal effects are strongest around W2 before gradually returning to BL_avg_ levels by W8. Despite absolute (ZT0-24), or lights-on (ZT0-12) and lights-off (ZT12-24) levels returning to normal, we have repeatedly observed time-of-day effects on W8. This suggests that certain structural changes to behavior are longer lasting than others, and recovery from oxycodone withdrawal may not be complete by W8. As such, we determined the magnitude and time-of-day dependence for slopes at each sleep stage on BL_avg_, W2, and W8 in 3-h bins (**Fig 7E**, NREM; **S5D Fig**, REM; **S6D Fig**, Wake; **right panels**). For NREM, HF slopes are significantly flattened during both lights-on and lights-off on W2 (**Fig 7E, middle panels**). When W2 slopes are calculated for 3-h bins, a more detailed picture emerges in which HF slopes are significantly increased for all but the ZT21-24 bin (**Fig 7E, right panel**), yet the most robust flattening of slopes clearly occurs at the beginning of lights-on. A 2-Way ANOVA reveals a 3-h bin × Treatment Day interaction (HF: F_(4.06, 40.6)_ = 7.171, p<0.0005), with Dunnett’s post hoc tests demonstrating significant pairwise differences in 1/f slopes between BL_avg_ and W2 for each time point except ZT21-24. For REM, there is a significant steepening of slopes (slopes become more negative) for both LF and HF EEG signals (**S5C, D Figs, middle panels**), but we noted that numerically, these effects are quite small. Therefore, it is fitting that quantification of LF and HF slopes from W2 in 3-h bins (**S5C, D Figs, right panels**) shows no significant differences between W2 and BLavg at any time point (LF: F(4.957, 48.16) = 1.859, ns; HF: F_(3.365, 32.69)_ = 1.965, ns). Finally, it is noteworthy that LF and HF slopes on W8 are not significantly different from those of BL_avg_ for any sleep stage, measured via CI, lights-on/off comparisons, or in 3-h bins.

This differs from the temporal effects of oxycodone withdrawal on % time in sleep stage (**Fig2D, E, F**) and NREM relative power (**Fig 6**).

### Chronic saline does not change sleep-wake rhythms

For these studies we used a within-subjects design in which each rat’s withdrawal data are compared to its average baseline data (BL_avg_). However, this design does not allow direct detection of any deterioration of EEG/EMG signal over time or possible effects of rat age, weight, and general health on EEG/EMG signal or sleep stage architecture and dynamics. To control for these possibilities, separate rats (N=3) were implanted with iPrecio pumps that infused saline for 14 days using conditions identical to those used for oxycodone (see **Fig. 1A**). Telemetry data were collected and analyzed in parallel with those from oxycodone rats. Because there are only 3 saline rats, experimental power for quantitative analysis is low. However, qualitative plotting of 3-h bin saline data for % time in each sleep stage, NREM relative power and LF/HF 1/f slopes across B4-B6 and W2, W4, W6, and W8 demonstrates a clear maintenance of the magnitude and temporal rhythmicity of sleep architecture and dynamics throughout the experiment (**Fig 8**). The one seeming outlier is NREM LF slopes (**Fig 8E, left panel)**. Upon close inspection, we see that the time-dependent slopes for W2, W4, W6, and W8 have the same shape and directionality as those for baseline, but they are shifted down such that the lowest 1/f slope value is -1.5 (W2-W8) rather than -1.0 (baseline). Regardless, the shapes and time courses of the saline NREM 1/f slopes at W2, W4, W6, and W8 are consistent across days, unlike those for the oxycodone withdrawal NREM 1/f slopes.

**Fig. 8:**
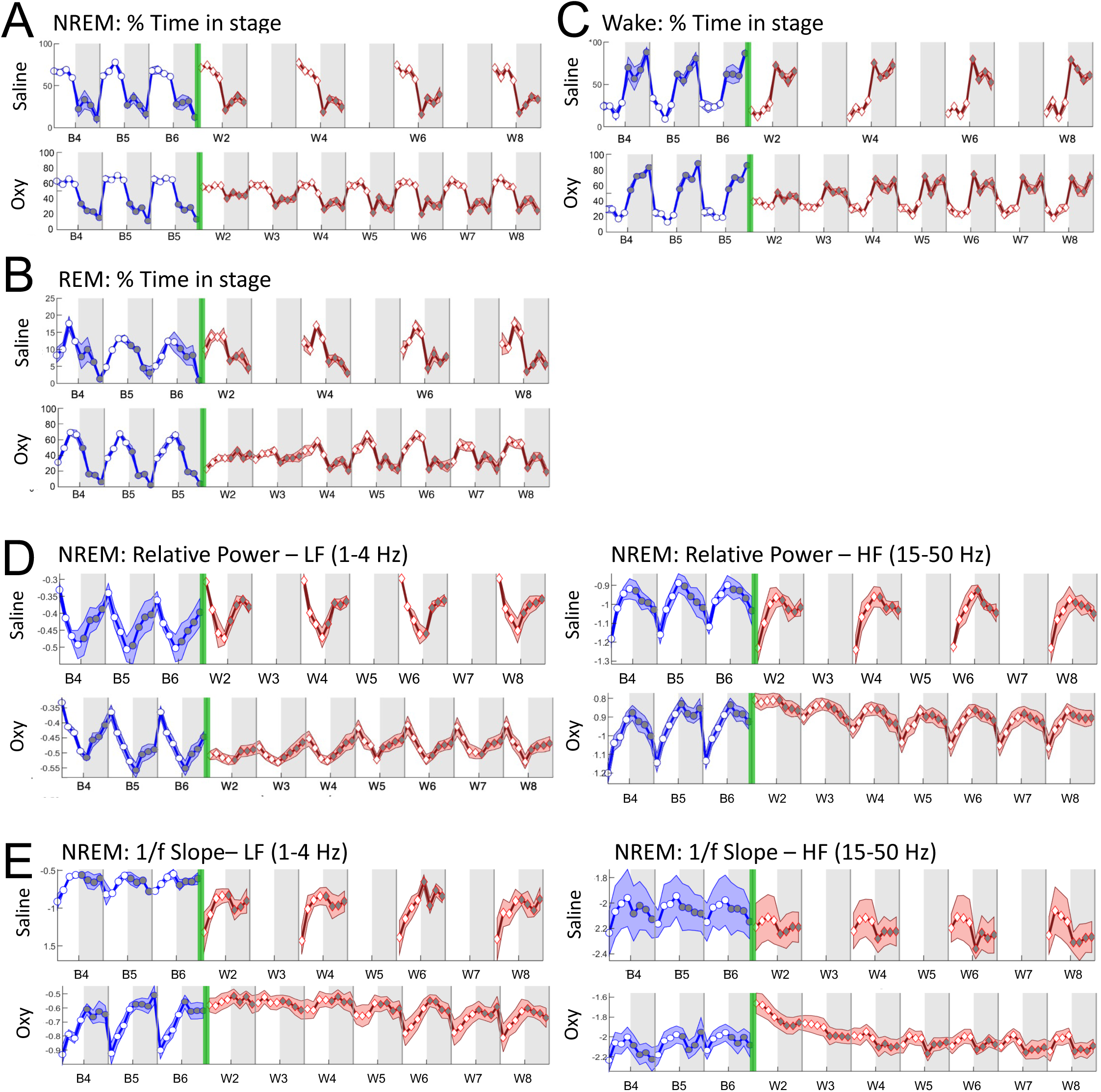
Chronic saline does not change sleep/wake rhythms. For control rats that received 14-d chronic saline infusions via iPrecio pumps (N=3), the mean % of total time (∓SEM) per 3-h bin spent in each sleep stage NREM, REM, and Wake is shown in the **top panels** of (**A, B**, and **C**, respectively) for baseline days 4, 5, and 6 (B4, B5, B6; blue line, circles) and for W2, W4, W6, and W8 (red line, diamonds). For comparison, the mean % of total time spent in each sleep stage in oxycodone-treated rats (N=11) undergoing spontaneous withdrawal (B4-B6 and W2-W8) is shown in the **bottom panels** of (**A, B**, and **C**). For control saline rats, the mean (∓SEM) LF (1-4 Hz) (**D, upper left panel**) and HF (15-50 Hz) (**D, upper right panel**) relative power per 3-h bin for B4, B5, B6 and W2, W4, W6, and W8 is shown. For comparison, the mean relative power for oxycodone-treated rats undergoing spontaneous withdrawal (B4-B6 and W2-W8) is shown in the **bottom panels** of (**D**). For control saline rats, the mean (∓SEM) LF (1-4 Hz) (**E, upper left panel**) and HF (15-50 Hz) (**E, upper right panel**) 1/f slope per 3-h bin for B4, B5, B6 and W2, W4, W6, and W8 is shown. For comparison, the mean 1/f slope for oxycodone-treated rats undergoing spontaneous withdrawal (B4-B6 and W2-W8) is shown in the **bottom panels** of (**E**). Underlying data is in **Fig8_Data.** *Abbreviations*: BL_avg_, Baseline average; W, withdrawal.

Even with the shifted NREM LF slopes discussed above, the temporal plots of sleep structure and dynamics during saline withdrawal remain almost identical to baseline plots. We conclude that 1) telemetry devices maintain reliable and consistent EEG/EMG recordings for at least the length of our experiments, and 2) the observed sleep disruptions measured during W2-W8 of spontaneous oxycodone withdrawal can be specifically attributed to oxycodone withdrawal itself.

## Discussion

In this study, we provide a comprehensive analysis of time-varying spontaneous oxycodone withdrawal effects on the circadian composition of discrete sleep stages and the dynamic spectral properties of the electroencephalogram (EEG) signal in male rats. Upon cessation of a 14-d non-contingent regimen of escalating-dose oxycodone, rats show a profound loss of pre-oxycodone (baseline) circadian rhythmicity of sleep/wake architecture (% time in NREM, REM, and Wake), temperature, and activity typically seen in nocturnal mammals. In addition, circadian-like rhythmicity of relative spectral band power (i.e., delta, alpha, beta and gamma) and LF aperiodic slope during NREM is dampened. During the first several days of spontaneous oxycodone withdrawal, rats spend less time asleep (NREM, REM) and more time awake during lights-on and more time asleep/less time awake during lights-off. This is broadly consistent with recent studies [19, 20] and with clinical observations of insomnia and daytime sleepiness in humans during opioid withdrawal and following chronic opioid use [6, 9, 10, 12]. Both sleep and wake states are more fragmented during early withdrawal, suggesting widespread instability across vigilance states. Finally, detailed analyses of EEG spectral power and aperiodic slope show the greatest oxycodone withdrawal effects on NREM sleep. Relative power shows a gradient of inversely proportional change with increasing frequency, which indicates a global change in the orientation of the EEG power spectrum—thus motivating a direct study of the dynamics of the EEG aperiodic slope. For NREM, oxycodone withdrawal results in a flattening of aperiodic slopes (slope magnitude closer to 0), broadly suggesting increased neural excitation [42, 44]. Together with the temporal dependence of sleep metrics on oxycodone withdrawal day, these foundational findings invite mechanistic studies to understand the neurobiological mechanisms engaged in opioid withdrawal’s disruption of sleep.

### Spontaneous oxycodone withdrawal disrupts the circadian rhythmicity and architecture of sleep

As previously reported, opioids such as oxycodone have acute effects on sleep processes [18]. In our study, the ZT0-2 subcutaneous infusion of high dose oxycodone (8 mg/kg) suppresses the circadian (baseline) increase in NREM and REM sleep and decrease in time awake that occurs when the lights turn on (ZT0). This effect lasts for ∼6 h after oxycodone infusion and is thought to result from both the infusion time course and predicted brain pharmacokinetics and pharmacodynamics of oxycodone [50]. Similarly, the ZT12-14 infusion of oxycodone suppresses the typical decrease in NREM and REM sleep and increase in time awake that occurs when the lights turn off (ZT12). During spontaneous withdrawal, however, sleep disruptions are robust, long-lasting, and not directly produced by drug infusion. These oxycodone withdrawal effects are the focus of this paper.

The circadian rhythmicity of NREM, REM, Wake, temperature, and activity under baseline conditions is clearly visualized in plots of the % time (**Fig. 2A, B, C**, blue lines) or of temperature and activity (**Fig. 3A, B**, blue lines) plotted in 3-h increments. Following baseline, the effects of spontaneous oxycodone withdrawal are shown for 7 days (W2-W8, red lines). For each output measure - except temperature - there is a transient flattening of circadian rhythmicity quantified using the circadian index (CI). The reduced CI (i.e., flattening) results from a decrease in the difference between daily maximum and minimum values of behaviors during both lights-on and lights-off. Temperature (subcutaneous measure) maintains its rhythmicity, but the magnitude of the typical lights-off increase is reduced during withdrawal. This dissociation between the flattening of circadian patterns in sleep behaviors and temperature is consistent with the finding that circadian variation in sleep stages is dissociated from body temperature [51]. Although spontaneous oxycodone withdrawal disrupts the % of time spent in each sleep/wake stage and temperature/activity compared to BL_avg_, the most robust differences occur primarily during lights-off. Indeed, even by W8, there are significant differences in sleep/wake times during lights-off but not lights-on. Further, withdrawal-induced disruptions of temperature and activity are only observed during lights-off.

Taken together, the temporal effects of spontaneous oxycodone withdrawal on daily rhythms and magnitudes of sleep/wake and temperature/activity indicate disruption of circadian rhythms. Although we did not test this directly, there is increasing evidence for chronic opioid administration and withdrawal to modulate molecular and behavioral circadian processes [52, 53]. For example, after 30 days of abstinence in human heroin users, disruptions in expression of PERIOD clock genes and diurnal rhythms of cortisol and endorphins persist [28]. Almost identical results were found in rats for up to 60 days of withdrawal from chronic morphine administration [27]. In the Li et al study (2010), blunted circadian expression of PERIOD genes occurred not just in the suprachiasmatic nucleus (SCN, central pacemaker of the brain), but also in the prefrontal cortex, nucleus accumbens, and amygdala - brain regions that regulate emotional states such as reward and aversion. Behaviorally, it has been shown that long-access heroin self-administration reversed the sleep-wake cycle in rats, and during abstinence, wake and NREM returned to baseline circadian rhythms, whereas REM sleep maintained its abnormalities for 3–6 days [54]. In addition to circadian processes, sleep regulatory mechanisms include homeostatic processes that track sleep need [29, 30, 31, 32], and allostatic processes such as stress [33]. Indeed, opioid withdrawal is an extreme stressor and considered an allostatic load [33], which is consistent with reports of insomnia from people with OUD during withdrawal [11, 55] To minimize experimental stress, we designed our experiments such that oxycodone administration and EEG/EMG recordings were wireless, constraining stress as much as possible to the experience of withdrawal itself.

### Sleep fragmentation during early oxycodone withdrawal disrupts depth of NREM sleep

NREM sleep is chiefly defined by the presence of low-frequency power associated with slow waves in the EEG spectrum [36]. Indeed, deep NREM is often referred to as slow wave sleep (SWS) [56], and low-frequency power is considered a surrogate of sleep depth. Under normal (baseline) conditions NREM low frequency power dissipates over the course of the sleep period (ZT0-12) and is indicative of a reduction in sleep pressure [29, 55, 57, 58]. In humans, NREM sleep is most commonly defined in three stages (N1-N3), with sleep depth going from low (N1) to high (N3) [59, 60, 61]. Recently, it has been shown that slow oscillation power itself is a reliable, objective, and continuous correlate of sleep depth [62].

In rodents, however, NREM is scored as a unitary stage, and the likely existence of NREM depth in rodents has been largely ignored. This is primarily due to rodents’ comparatively short and rapid fluctuation between NREM and REM sleep, with cycles lasting around 10 min in rodents vs 1-2 hours in humans [63, 64]. Sleep spindle activity, the hallmark of N2 sleep in humans, is spectrally diffuse in rodents [65], making it harder to define, detect, and score. The convention of a single NREM state in rodents is likely more a feature of practical scoring logistics, rather than a mechanistic statement on constant sleep depth. Thus, despite lack of standardized differentiation between NREM stages, it would not be impertinent to discuss changes in low-frequency power as being related to depth of NREM sleep in rodents.

Figure 5 shows circadian-like periodicity throughout baseline days in all NREM EEG frequency bands. In particular, we observe NREM delta power (Fig. 5A) to be at its highest at ZT0 (lights on), linearly decrease from ZT0-12, and then increase again once lights are off, a pattern consistent with dissipation of sleep pressure and changing sleep depth. Conversely, we see the opposite pattern for frequencies higher than 8 Hz, which are typically associated with wakefulness and arousal [34] (Fig. 5C**-E**). The reduction in delta power during early oxycodone withdrawal is consistent with a recent study in mice [19] showing decreased delta power during morphine withdrawal, which was hypothesized to represent a reduction in sleep pressure. However, we see a significant but small (∼5%) decrease in lights-on % time in NREM during W2-W5, with a large (∼20%) significant increase during lights-off for W2-W3 (Fig. 2D middle panel). Consequently, the observed decrease in delta power is not likely principally driven by major reductions in sleep pressure. Rather, we observe a significant increase in the number of lights-on NREM bouts on W2 and a significant decrease in bout duration W2-W6 (Fig. 4B, left panel). Specifically, lights-on NREM W2 shows a ∼50% increase in the number of bouts and a ∼25% decrease in bout duration, suggesting strong fragmentation of NREM sleep during early withdrawal. Moreover, during lights-off, there is a significant increase in the number of bouts for W2-W4, with no significant change in duration, consistent with high sleep pressure leading to extended sleep related to recovery after sleep disruption. These findings are therefore consistent with a scenario in which fragmentation prevents deep NREM sleep, rather than one in which sleep pressure is low.

NREM fragmentation has varied effects on low-frequency power, as evidenced by conflicting reports from the narcolepsy and obstructive sleep apnea literature [66, 67, 68]. In the context of opioid withdrawal in humans, it is possible that NREM fragmentation increases low-frequency power, and hence sleep pressure, to compensate for reduced sleep. On the other hand, NREM fragmentation could also decrease low-frequency power such that the subject does not have sufficient time to enter deep NREM sleep. In general, the evidence suggests that sleep fragmentation plays an important role in the changes observed during early withdrawal. Finally, it cannot be ignored that opioid withdrawal is an extreme stressor for the animal, thus providing a strong allostatic load on sleep regulation [69, 70].

Taken together, our data support the utility of viewing NREM sleep as a continuum across sleep depth (as in humans), rather than a unitary stage. However, rhythmic changes in NREM sleep depth are not the sole driver of the amount of time spent in sleep stages under baseline conditions or after oxycodone withdrawal. As mentioned earlier, circadian effects likely play an important role: even with withdrawal-induced flattening of circadian periodicities, clear changes in sleep metrics are consistently observed at lights-on and lights-off.

### Early withdrawal disrupts the aperiodic component of the EEG

While it is tempting to view changes in each EEG frequency band in isolation, it is important to consider the EEG spectrum as a whole. When looking at the NREM EEG power bands together during baseline (Fig. 5), the results indicate coordinated, global changes in the overall slope of the EEG spectrum with a pivot point around theta, such that low and high frequencies are anti-correlated. Thus, the within-NREM periodicity can be said to predominantly reflect cyclical changes in the aperiodic component of the EEG signal. This is confirmed through visualization of the mean NREM relative power spectra across conditions (**Fig 7A**), which shows a tilt in the slope of the spectrum pivoting in the theta range on W2. This is directly quantified through direct estimation of low (1-4Hz) and high (15-50Hz) frequency (LF, HF, respectively) components of the NREM 1/f slope across the experiment (Fig. 7D, E). Upon withdrawal, both LF and HF components show a significant (LF: W2-W5, HF: W2-W3) flattening (1/f slopes approach 0) for lights-on with marked loss of periodicity (**Fig 7B**, LF; **Fig 7C**, HF). Given the duration and significance of the difference, LF lights-on slope (**Fig 7D**, middle panel) is the most sensitive indicator of withdrawal studied herein, making this metric a strong candidate for a potential EEG biomarker.

Mechanistically, changes in the aperiodic slope have been hypothesized to reflect alterations in the excitatory-inhibitory balance (EI) of the brain. Indeed, pharmacological excitation of neural activity is associated with flatter slopes (i.e., slope values closer to 0), whereas inhibition is associated with steeper slopes [40, 42, 44]. Additionally, flatter 1/f slopes during NREM can also indicate a decrease in inhibitory receptor density [43, 71]. A caveat with interpreting scaling changes in 1/f slopes as shifts in EI balance is that, to date, the referenced pharmacological studies above often used compounds with multiple dose- and brain region-dependent targets. Despite this caveat, it has been clearly demonstrated that 1/f activity is robustly modified by drugs, cognitive tasks, and psychiatric conditions [72, 73]. Taken together, our finding that spontaneous oxycodone withdrawal flattens low frequency NREM aperiodic slopes is commensurate with the idea that EI balance is shifted towards greater excitability in neural circuits associated with sleep.

Studies on the molecular and electrophysiological [29, 30, 31, 32, 33] actions of chronic opioid exposure and withdrawal have repeatedly shown a rebound increase in neural excitation once MOR activation ceases with drug withdrawal [23, 24, 25]. The functional consequences of rebound excitation are brain region-dependent, with several MOR-expressing brain regions contained within sleep circuits [22]. As one example, morphine withdrawal increases expression of the immediate early gene, cFos, in the region of the lateral hypothalamus that produces the wake-promoting neuropeptide orexin [74]. Orexin receptor antagonists have been shown to decrease opioid withdrawal symptoms, opioid tolerance, and motivation to seek opioids in rodents, suggesting the peptide itself plays an important role in the allostatic response to chronic opioid exposure [75, 76]. The dual orexin receptor antagonist, suvorexant, has FDA approval for the treatment of insomnia [77] and is in clinical trials for treatment of opioid withdrawal sleep disruptions [78].

In summary, these foundational studies use principled technical and computational methods to characterize how, and for how long, oxycodone withdrawal impacts sleep structure and dynamics in rats. Despite the growing focus of clinical research on sleep disturbance in people receiving medication for opioid use disorder, there are very few human research studies thoroughly examining how opioid withdrawal impacts sleep, and there have been only a few preclinical studies focused on opioid withdrawal and sleep; notably two of them recent studies [19, 20]. This is a massive gap in knowledge, as sleep disruption is both common and persistent among people receiving medication for opioid use disorder [14,78]. The few studies examining treatment of insomnia for opioid use disorder have shown limited efficacy for common sleep medications [79, 80] highlighting the need for improved mechanistic understanding of the impact of opioids of sleep link to identify better treatment targets for alleviating sleep disruption in people with opioid use disorder. We believe this in-depth characterization of several translatable sleep measurements, including basic sleep architecture, frequency band-dependent oscillatory activity. and aperiodic EEG activity will allow fundamental questions about the mechanisms underlying opioid withdrawal-induced sleep deficits to be answered and advance treatment development. In addition, it is likely that employing these methods clinically will enable sleep metrics to serve as biomarkers for the magnitude and time course of the oxycodone withdrawal syndrome in humans.

## Materials and Methods

### Animals

Adult male Sprague Dawley rats (Charles River Laboratories, Wilmington, MA]) weighing 250-275g upon arrival were initially group-housed in the animal care facility. After surgery, rats were individually housed in a satellite “sleep room” designated specifically for rodent sleep studies, with each cohort of rats being the sole occupants of the sleep room. Both the animal care vivarium and sleep room were kept on a 12h light/dark (7:00 AM lights on / 7:00 PM lights off) cycles. Standard rat chow (Lab Diet 5012) and water were available *ad libitum*. To ensure reproducibility of our data, 5 independent cohorts of rats were tested (Total=14 rats). Cohort 1 (N=3), Cohort 2 (N=3), Cohort 3 (N=1), Cohort 4 (N=3), Cohort 5 (N=1). The housing and treatment of rats were approved by the McLean Hospital Institutional Animal Care and Use Committee and followed guidelines set by the National Institutes of Health.

### Surgeries

At least one week after arrival, rats underwent surgery (body weight at time of surgery was 310-330g). Briefly, rats were anesthetized with isoflurane (Covetrus; isoflurane at 4-5% for induction, 1.5-3% for anesthesia maintenance; 1.5L/min O_2_ flow) and subcutaneously implanted with both a telemetry device [HD-S02; Data Sciences International (DSI)] and a programmable infusion pump [iPrecio SMP-200; DURECT Corporation (iPrecio pump)].

*Telemetry surgeries*: anesthetized rats were implanted with a telemetry device (HD-S02; DSI, USA). Telemetry devices contain 4 biopotential leads: 2 serve as EMG electrodes and 2 as EEG electrodes. For each implant, the EMG electrode leads were threaded through the cervical trapezius muscle via a small incision made with a 20-gauge needle and was anchored to the muscle using non-dissolvable silk sutures. The EEG electrode leads were individually secured to two skull screws (Plastics One, cat #00-80 x 1/16; 1.6mm long) that contacted dura. The negative EEG lead was secured to the screw at +2.0mm anteroposterior and at lateral left 1.5mm, and the positive EEG lead was secured to the screw at -7.0mm anteroposterior and lateral right -1.5mm – all relative to Bregma. The screw plus lead assemblies were held in place on the skull with dental cement (Ortho-Jet Package; Lang Dental).

*iPrecio drug pump surgeries*: Immediately after implanting the telemetry device, iPrecio drug pumps were subcutaneously implanted. The pump tubing, from which drug is released, was cut to 2.5-3.0cm prior to implantation. A subcutaneous pocket was created at 3-4cm posterior to the scapula and 2cm lateral and parallel to the spine, and the pre-programmed iPrecio pump was inserted. To secure the pump within the pocket, a suture was threaded through the underlying external oblique muscle and then through a suture anchor located on the pump. After surgery, rats were individually housed in the sleep room.

### Physiological telemetry recordings

Four to five days after surgery, telemetry devices were activated by a magnet (DSI), which puts the devices on “standby”, allowing control via DSI Ponemah software. Receivers (RPC-1 PhysioTel; DSI) that detect radio signal are placed underneath each rat cage to detect AM signals emitted by the telemetry transmitters. Data was recorded 24 h/d throughout the experiment (see Fig. 1A) and included 500-Hz interpolated sampling of EEG and EMG and 10-s bins of mean subcutaneous temperature and activity counts. Zeitgeber time 0 (ZT0) was defined as 7:00AM. EEG, EMG, motor activity, and temperature were analyzed using Neuroscore (DSI).

### Escalating-dose oxycodone regimen to induce dependence

Prior to implantation, iPrecio pumps were programmed using the iPrecio Management System and Software (IMS-200; ALZET) to infuse either saline or oxycodone twice a day for two hours each (ZT0-2 and ZT12-14). For the first 9-12 days after surgery, the pumps infused 0.9% saline at 15 μl/h for 2h. Figure 1A shows a schematic of the experimental design. Telemetry data from the last 3 days of saline infusions were used as baseline. Following baseline, saline was removed from the pumps and replaced with oxycodone (0.5 mg/kg). Separate rats received saline throughout the 14-d pump delivery regimen and served as controls for any effects of surgeries, interoceptive effects of infusions, isoflurane exposures, aging/weight gain on sleep. In addition, saline rats controlled for telemetry implant and receiver stability over the course of the experiment. The iPrecio pumps were emptied and refilled with the appropriate concentration of oxycodone (or saline) 4 times, always between ZT7-12. To refill the pumps, rats were briefly anesthetized with 4-5% isoflurane, and a 26-gauge syringe was used to withdraw all fluid in the reservoir and add new drug via a port on the pump that could be felt through the skin. The first oxycodone infusion was programmed for ZT12-14 on day 1 and the last for ZT0-2 on withdrawal day 1 (W1), such that rats received a total of 28 oxycodone (or saline) infusions over 14 days. The first 4 infusions were delivered at a rate of 15ul/h, and the remaining infusions were delivered at a rate of 30μl/h. To demonstrate that the escalating-dose oxycodone regimen produced dependence, rats were injected with the opioid receptor antagonist, naloxone (1.0 mg/kg, SC), or saline 30-45 min after the last oxycodone (8.0 mg/kg) infusion ended to assess naloxone-precipitated somatic withdrawal signs. Immediately after naloxone (or saline) injections, rats were videotaped in their home cages for at least 40 minutes, and the videos subsequently scored for withdrawal signs by a trained experimenter, as previously described [25]. Because rats received either naloxone or saline injection on W1, we did not consider sleep data collected on that day to cleanly represent spontaneous withdrawal. Rather, ZT0 on W2 marked the start of spontaneous oxycodone withdrawal in our study.

### Sleep scoring

Using a set of rules based on a combination of behavior and EEG events, sleep states were categorized into wake, rapid eye movement (REM) sleep, and non-REM (NREM) sleep [47]. Sleep scoring was performed manually by a trained experimenter using Neuroscore, in which 10-s epochs of EEG/EMG waveforms were used to score sleep states using standard methods [47]. Sleep scoring was verified at random epochs by a second trained experimenter to confirm inter-rater consistency.

### Circadian metrics

Percent time in Wake, NREM and REM as well as activity counts and temperature were computed for 3 baseline (BL) days and for withdrawal days 2-8 (W2-W8). During the first several days of oxycodone withdrawal, there was a clear, qualitative reduction in the nocturnal amplitudes of our behavioral measures: Wake, NREM, REM, activity, and temperature. Therefore, we computed the circadian index (CI) for BL and W2-W8 for each measure. We defined the CI for a given measure as the difference between the maximum 3h bin value and the minimum 3h bin value on a particular day relative to each individual rat’s average baseline difference.

### Spectral estimation

To characterize changes in neural activity during withdrawal, we performed spectral estimation on the EEG data, which provides a basis for quantifying the frequency content of the EEG signal. Spectral estimation was performed using the multitaper approach, which is an estimator with optimized bias and variance properties [81] commonly used for sleep EEG analyses [36]. Multitaper spectrograms were computed with the following parameters: frequency range: 0.5 - 50Hz, time-bandwidth product: 5, number of tapers: 9, window size: 10s, window step: 5s. For analyses of specific conditions or time segments, fixed spectra were computed by averaging the spectrogram over the given period. See Prerau et. al 2017 [36] for a detailed discussion of multitaper parameter selection. To adjust for inter-rat variability in recording, we computed relative power, defined as the power within each frequency bin divided by the total power between 0.5 and 50Hz, which was log-transformed for analysis.

### Parameterizing the aperiodic slope

While numerous sophisticated approaches have been proposed to model the non-linear aspects of the aperiodic EEG spectrum [41, 71], the most straightforward method transforms the spectrum into the log-log space, in which the decay constant can be parameterized as the slope of a line. This linear approach has been shown to have negligible deviation in population estimates relative to the non-linear approaches [41]. We fit the piecewise low (1-4Hz) and high (15-50Hz) components of the log-log transformed spectra using least-squares linear regression with an offset, from which the value of the slope parameters was used as the estimate. It should be noted that there is no scientific consensus for the frequency range of the high and low components, which are determined purely ad-hoc based on the application and whether human or animal models are being used [42, 43, 44].

### Experimental Design and Statistical Analyses

Statistical comparisons were performed using GraphPad Prism version 10 (GraphPad Software). For all telemetry data, we used a within-subjects design in which each rat’s withdrawal data was compared to the average of data from baseline days 4, 5, and 6 (see Schematic, Fig.1). We considered spontaneous withdrawal to start at ZT0 on W2, because this was the first full withdrawal day not confounded by additional drug treatments (i.e. on W1, rats received their last oxycodone infusions from ZT0-2 followed by either an acute injection of naloxone or saline at ZT0.5). For time course data in which % Time, CI, bout #, bout length, relative power, or 1/f slopes are compared to BL_avg_, normality tests were conducted. These data tended not to meet requirements for normal distribution due to one cohort of 3 rats whose sleep stage and temperature, but not spectral, data were consistently shifted and the cause of non-normality. All data were included in analyses, as there was no cause to exclude them. To be conservative in our analyses, we used the nonparametric Friedman’s Test on these data, followed by Dunn’s multiple comparison tests. For 2-way analyses in which 3-h time bins of % Time, relative power, or 1/f slopes from W2, W8, and BL_avg_ were compared, normality tests (e.g., D’Aostino and Pearson, Anderson-Darling tests), which were fairly evenly split between normal and non-normal distributions. Given the lack of a reliable non-parametric test for 2-Way ANOVA, and the similarity of statistical results for other data when tested with both parametric and non-parametric tests, we used 2-way, repeated measures ANOVA followed by Dunnett’s multiple comparison tests. Significance level was set at *p*<0.05.

