## Supplementary Figures for "Spontaneous oxycodone withdrawal disrupts sleep, circadian, and electrophysiological dynamics in rats"

**Supplementary Material**
**Supplementary Figure 1**

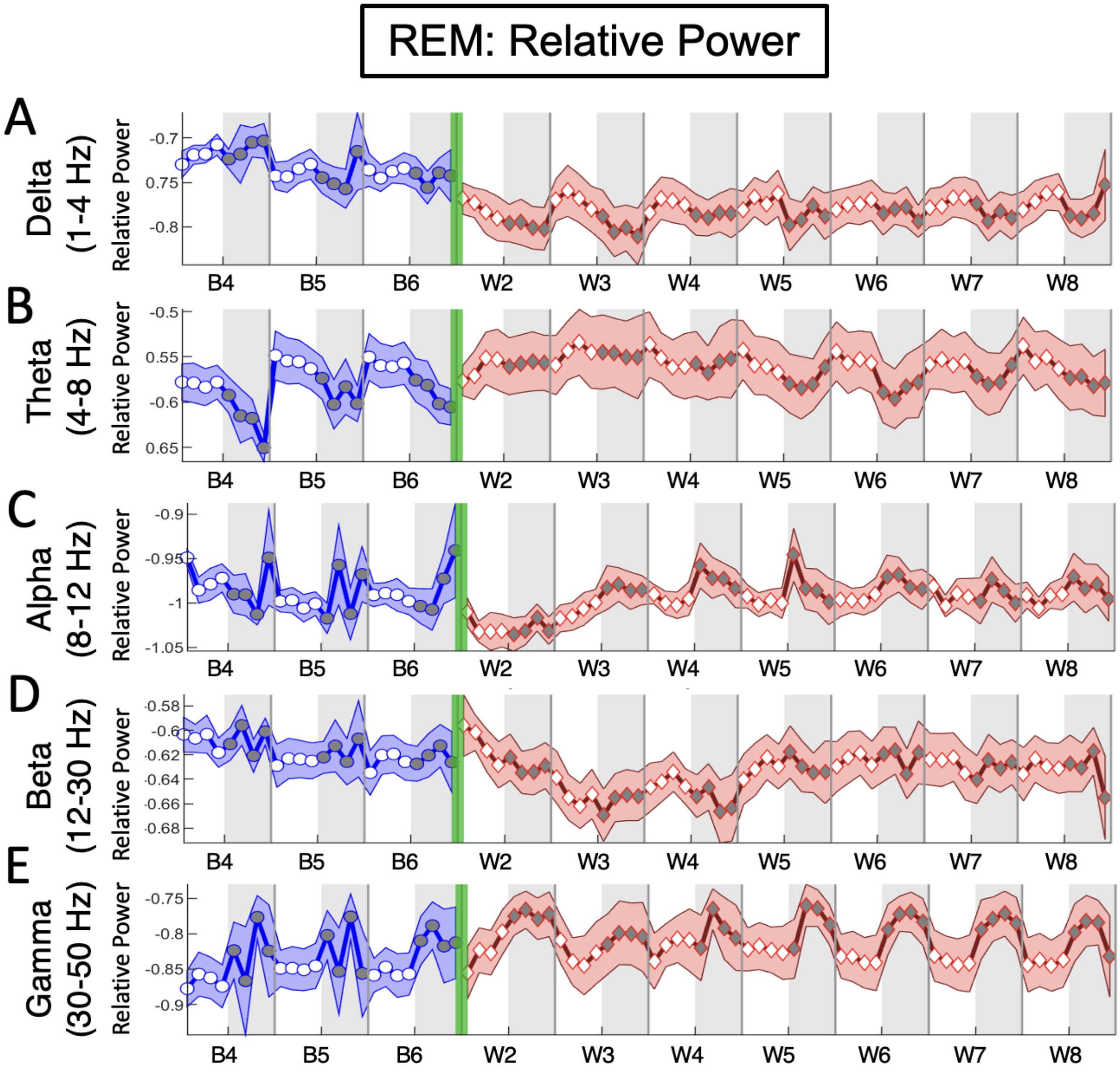


**S1 Fig: REM relative power plotted continuously throughout baseline and oxycodone withdrawal for multiple frequency bands shows little change.**
The mean relative power for REM is plotted in 3-h bins (∓SEM) for (**A**) Delta (1-4Hz), (**B**) Theta (4-8Hz), (**C**) Alpha (8-12Hz), (**D**) Beta (12-30Hz) and (**E**) Gamma (30-50Hz) frequency bands throughout baseline days 4, 5, and 6 (B4, B5, B6; blue line, circles) and spontaneous withdrawal days 2-8 (W2-W8; red line, diamonds). Vertical green bars between B6 and W2 represent the 14-d escalating oxycodone dose regimen and W1. Grey shaded regions represent ZT12-24 (lights off), and unshaded regions represent ZT0-12 (lights on). N=11 rats. **Underlying data is in "S1 Fig_Data".**

**Supplementary Figure 2**

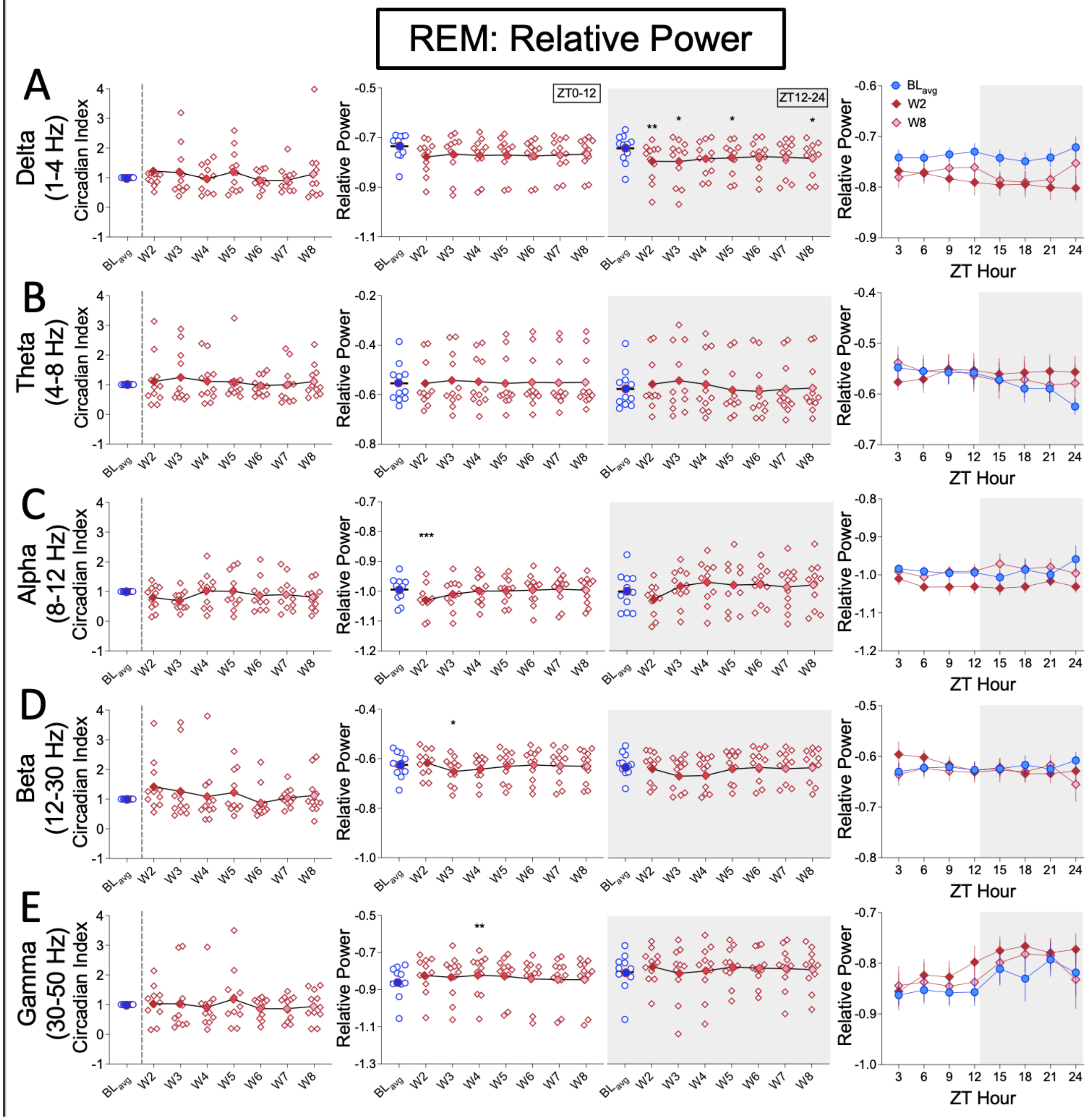


**S2 Fig: Spontaneous oxycodone withdrawal has minimal impact on REM power rhythms.**
For relative power during REM, the mean (∓SEM) Circadian Index (CI; **left panels**), mean (∓SEM) relative power per frequency band for ZT0-12 or ZT12-24; **center panels**, and the mean (∓SEM) relative power per frequency band in 3-h bins for

$BL_{avg}$, W2, and W8 (**right panels**) are shown for (**A**) Delta 1-4Hz, (**B**) Theta (4-8Hz), (**C**) Alpha (8-12Hz) (**D**) Beta (12-30Hz), (**E**) Gamma (30-50Hz) frequency bands. Non-parametric Friedman with Dunn's post hoc tests (**left and center panels**), and two-way repeated measures ANOVA with Dunnett's post hoc tests (**right panels**) compared W2-W8 to $BL_{avg}$. ***$p<0.001$.**$p<0.01$, *$p<0.05$. N=11 rats. **Underlying data is in "S2 Fig_Data".** *Abbreviations*: ZT, zeitgeber time; $BL_{avg}$, Baseline average; W, withdrawal.

**Supplementary Figure 3**

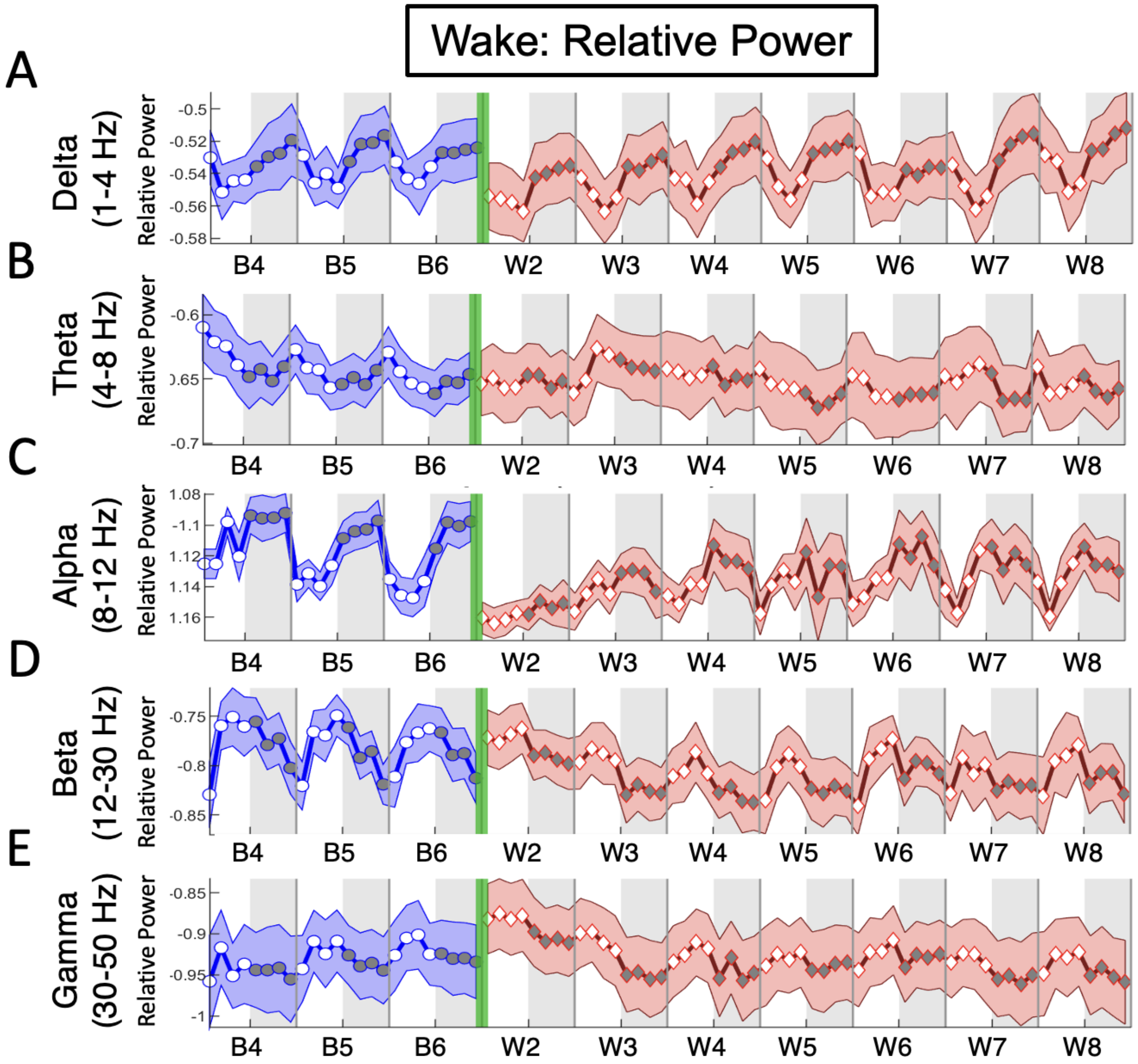


**S3 Fig: Wake relative power plotted continuously throughout baseline and oxycodone withdrawal for multiple frequency bands shows little change.**
The mean relative power for Wake is plotted in 3-h bins (∓SEM) for (**A**) Delta (1-4Hz), (**B**) Theta (4-8Hz), (**C**) Alpha (8-12Hz), (**D**) Beta (12-30Hz) and (**E**) Gamma (30-50Hz) frequency bands throughout baseline days 4, 5, and 6 (B4, B5, B6; blue line, circles) and spontaneous withdrawal days 2-8 (W2-W8; red line, diamonds). Vertical green bars between B6 and W2 represent the 14-d escalating oxycodone dose regimen and W1. Grey shaded regions represent ZT12-24 (lights off), and unshaded regions represent ZT0-12 (lights on). N=11 rats. **Underlying data is in "S3 Fig_Data".**

**Supplementary Figure 4**

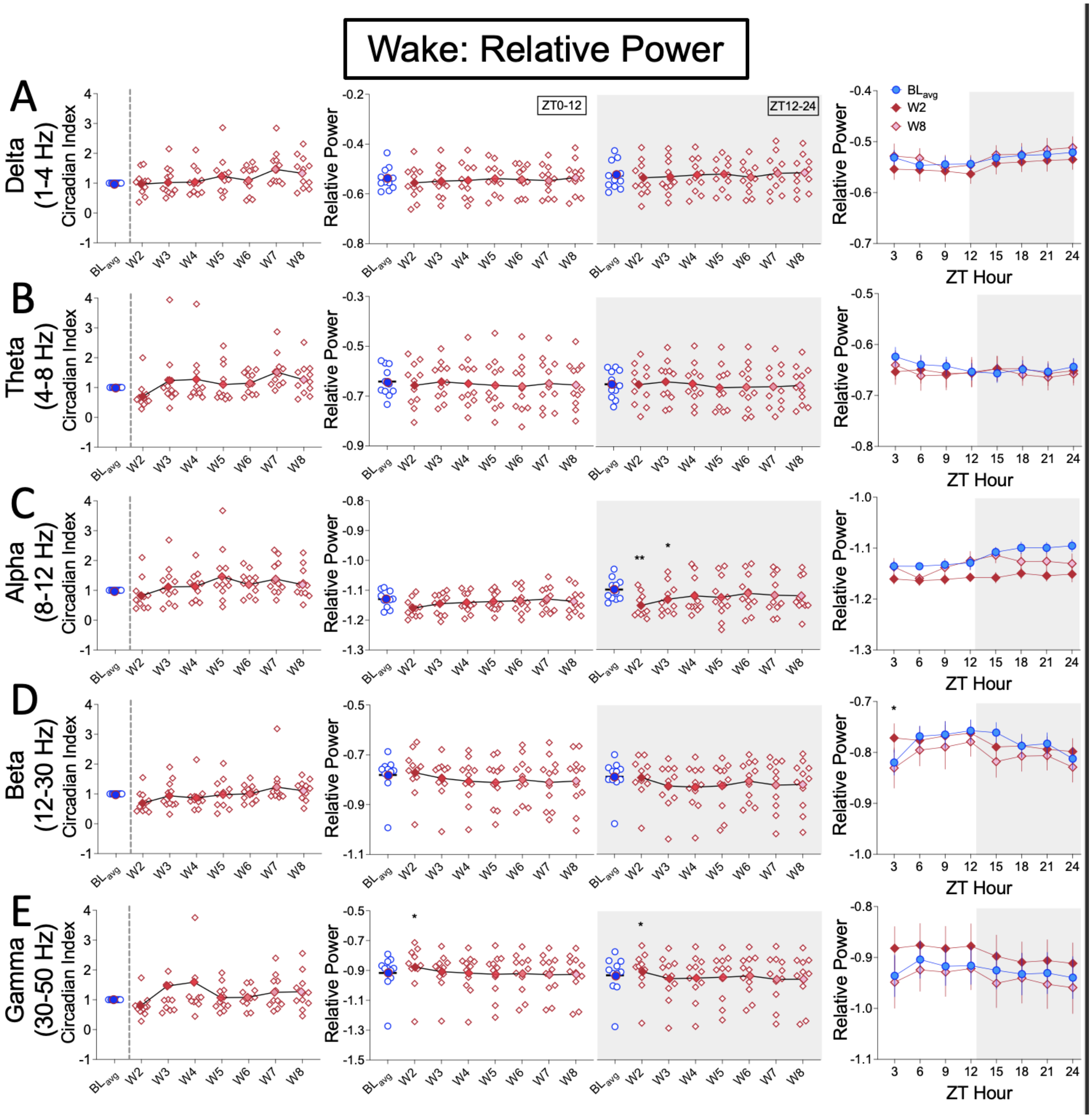


**S4 Fig: Spontaneous oxycodone withdrawal has minimal impact on Wake power rhythms.**
For relative power during Wake, the mean (∓SEM) Circadian Index (CI; **left panels**), mean (∓SEM) relative power per frequency band for ZT0-12 or ZT12-24; **center panels**, and the mean (∓SEM) relative power per frequency band in 3-h bins

for $BL_{avg}$, W2, and W8 (**right panels**) are shown for (**A**) Delta 1-4Hz, (**B**) Theta (4-8Hz), (**C**) Alpha (8-12Hz) (**D**) Beta (12-30Hz), (**E**) Gamma (30-50Hz) frequency bands. Non-parametric Friedman with Dunn's post hoc tests (**left and center panels**), and two-way repeated measures ANOVA with Dunnett's post hoc tests (**right panels**) compared W2-W8 to $BL_{avg}$. $^{**}p<0.01$, $^{*}p<0.05$. N=11 rats. **Underlying data is in "S4 Fig_Data".** *Abbreviations*: ZT, zeitgeber time; $BL_{avg}$, Baseline average; W, withdrawal.

**Supplementary Figure 5**

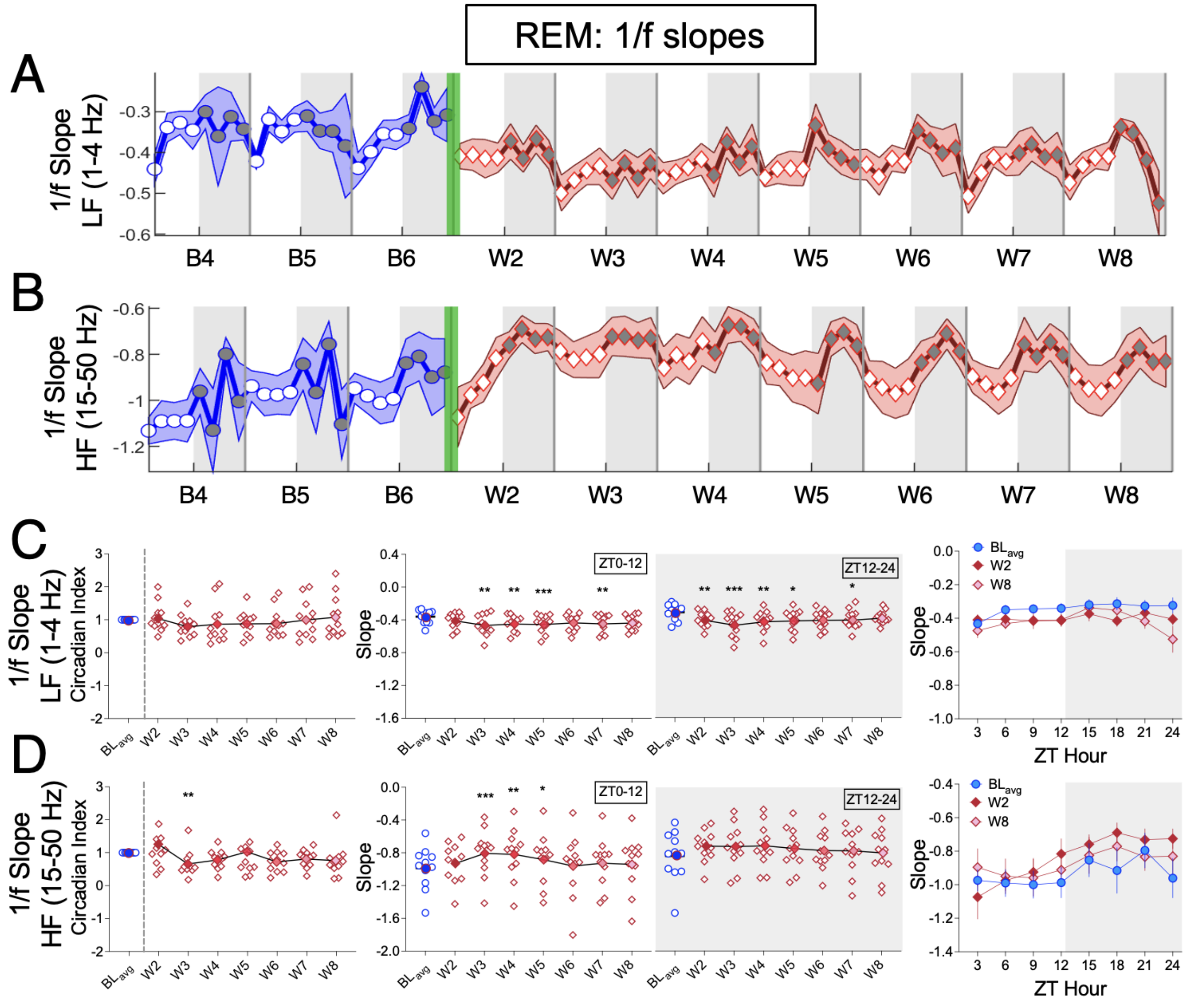


**S5 Fig: Spontaneous oxycodone withdrawal increases REM 1/f slopes in the low frequency range and increases slopes in the high frequency range in a withdrawal day-dependent manner.**
The mean REM 1/f slope (∓SEM) per 3−h bin is plotted for (**A**) LF (1-4 Hz) and (**B**) HF (15-50 Hz) ranges across baseline days 4, 5, and 6 (B4, B5, B6; blue line, circles) and spontaneous withdrawal days 2-8 (W2-W8; red line, diamonds). Vertical green bars between B6 and W2 represent the 14-d escalating oxycodone regimen and W1. For both LF (**C**) and HF (**D**) REM slopes, the mean (∓SEM; including individual rat values) Circadian Index (**left panels**), 1/f slopes (ZT0-12 or ZT12-24; **center panels**), and 3-h bin data for $BL_{avg}$, W2, and W8 (**right panels**) are shown. Non-parametric Friedman tests with Dunn's post hoc tests (**left and center panels**), and two-way repeated measures ANOVA with Dunnett's post hoc tests (**right panels**) compared W2-W8 to $B_{avg}$. ***p<0.001.**p<0.01, *p<0.05. N=11 rats. **Underlying data is in "S5 Fig_Data".** *Abbreviations*: ZT, zeitgeber time; $BL_{avg}$, Baseline average; W, withdrawal.

**Supplementary Figure 6**

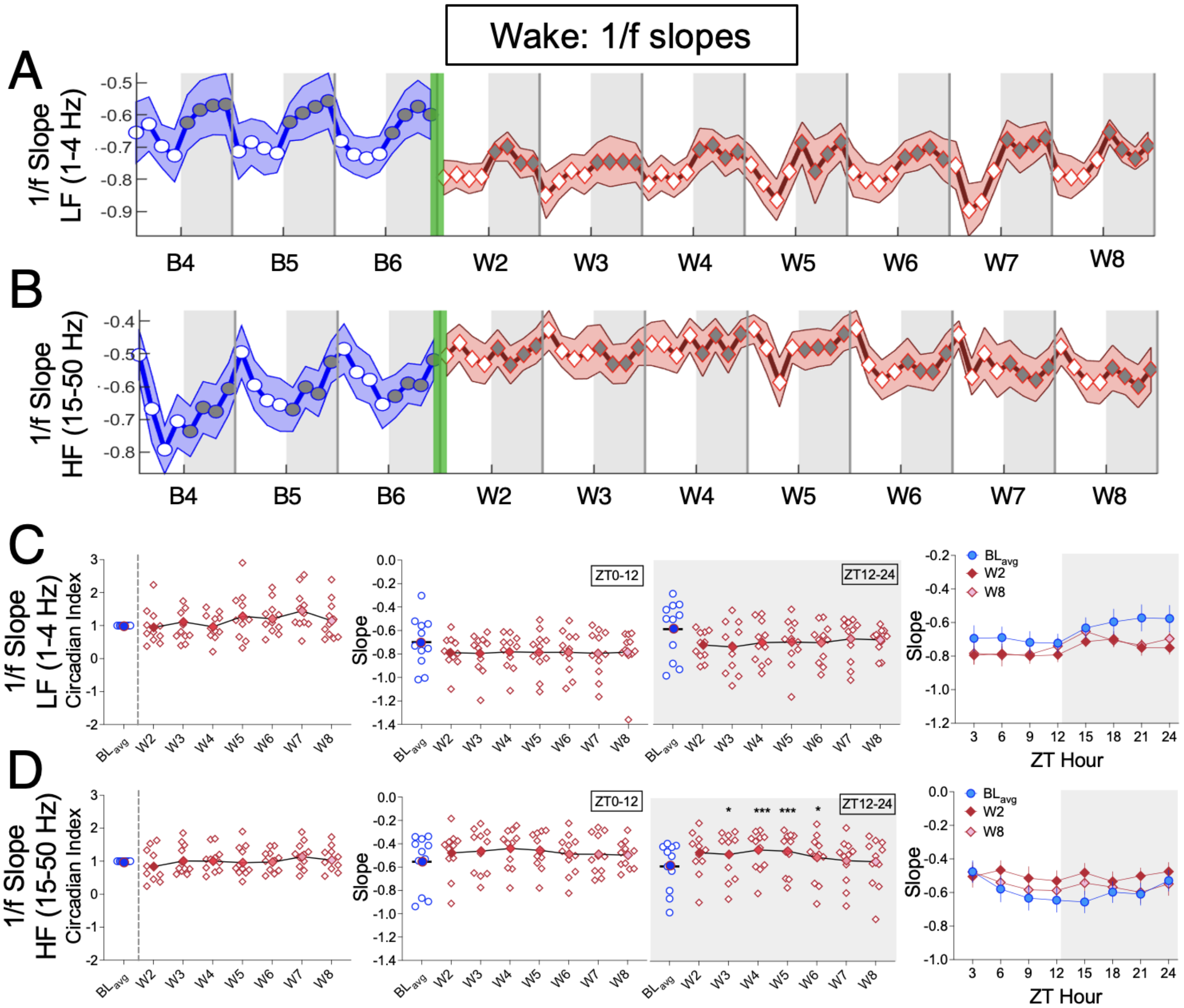


**S6 Fig: Spontaneous oxycodone withdrawal flattens Wake 1/f slopes in the high frequency range during lights-off in a withdrawal day-dependent manner.**
The mean Wake 1/f slope (∓SEM) per 3–h bin is plotted for (**A**) LF (1-4 Hz) and (**B**) HF (15-50 Hz) ranges across baseline days 4, 5, and 6 (B4, B5, B6; blue line, circles) and spontaneous withdrawal days 2-8 (W2-W8; red line, diamonds). Vertical green bars between B6 and W2 represent the 14-d escalating oxycodone regimen and W1. For both LF (**C**) and HF (**D**) Wake slopes, the mean (∓SEM; including individual rat values) Circadian Index (**left panels**), 1/f slopes (ZT0-12 or ZT12-24; **center panels**), and 3-h bin data for $BL_{avg}$, W2, and W8 (**right panels**) are shown. Non-parametric Friedman tests with Dunn's post hoc tests (**left and center panels**), and two-way repeated measures ANOVA with Dunnett's post hoc tests (**right panels**) compared W2-W8 to $BL_{avg}$. ***p<0.001, *p<0.05. N=11 rats. **Underlying data is in "S6 Fig_Data".** *Abbreviations*: ZT, zeitgeber time; $B_{avg}$, Baseline average; W, withdrawal
